# Surface-modified measles vaccines encoding oligomeric, fusion-stabilized SARS-CoV-2 spike glycoproteins bypass measles seropositivity, boosting neutralizing antibody responses to omicron and historical variants

**DOI:** 10.1101/2022.12.16.520799

**Authors:** Miguel Á. Muñoz-Alía, Rebecca A. Nace, Baskar Balakrishnan, Lianwen Zhang, Nandakumar Packiriswamy, Gagandeep Singh, Prajakta Warang, Ignacio Mena, Riya Narjari, Rianna Vandergaast, Adolfo García-Sastre, Michael Schotsaert, Stephen J. Russell

## Abstract

Serum titers of SARS-CoV-2 neutralizing antibodies (nAb) correlate well with protection from symptomatic COVID-19, but decay rapidly in the months following vaccination or infection. In contrast, measles-protective nAb titers are life-long after measles vaccination, possibly due to persistence of the live-attenuated virus in lymphoid tissues. We therefore sought to generate a live recombinant measles vaccine capable of driving high SARS-CoV-2 nAb responses. Since previous clinical testing of a live measles vaccine encoding a SARS-CoV-2 spike glycoprotein resulted in suboptimal anti-spike antibody titers, our new vectors were designed to encode prefusion-stabilized SARS-CoV-2 spike glycoproteins, trimerized via an inserted peptide domain and displayed on a dodecahedral miniferritin scaffold. Additionally, to circumvent the blunting of vaccine efficacy by preformed anti-measles antibodies, we extensively modified the measles surface glycoproteins. Comprehensive *in vivo* mouse testing demonstrated potent induction of high titer nAb in measles-immune mice and confirmed the significant incremental contributions to overall potency afforded by prefusion stabilization, trimerization, and miniferritin-display of the SARS-CoV-2 spike glycoprotein, and vaccine resurfacing. In animals primed and boosted with a MeV vaccine encoding the ancestral SARS-CoV-2 spike, high titer nAb responses against ancestral virus strains were only weakly cross-reactive with the omicron variant. However, in primed animals that were boosted with a MeV vaccine encoding the omicron BA.1 spike, antibody titers to both ancestral and omicron strains were robustly elevated and the passive transfer of serum from these animals protected K18-ACE2 mice from infection and morbidity after exposure to BA.1 and WA1/2020 strains. Our results demonstrate that antigen engineering can enable the development of potent measles-based SARS-CoV-2 vaccine candidates.

## Introduction

For the second year since severe acute respiratory syndrome coronavirus (SARS-CoV-2) was first identified, coronavirus diseases 19 (COVID-19) ranked as the third leading cause of death after heart disease and cancer ^1^. The number of lives taken globally by COVID-19 now exceed more than 6.5 million, and the number of cases is above 605 million. The pandemic has disrupted lives across the globe and triggered the deepest recession since World War II ^2^. Even though highly immunogenic and efficacious COVID-19 vaccines have been deployed, the continual emergence of immune-evasive variants of SARS-CoV-2 combined with the waning efficacy of SARS-CoV-2 vaccines still represents a major global health challenge ^3–6^.

Similar to other coronavirus infections, SARS-CoV-2 infection is mediated by homotrimeric class-I membrane-bound viral spike (S) proteins, which comprise an S1 domain containing the receptor-binding domain (RBD) that mediates attachment to the host cell, as well as an S2 domain containing the fusion peptide that initiates fusion with the host cell membrane ^7, 8^. Binding to the receptor carboxypeptidase angiotensin-converting enzyme 2 (ACE2) and proteolytic cleavage are thought to trigger the dissociation of the S1 domain and irreversible refolding of the S2 domain, leading to fusion between the viral and cellular membranes for cell entry ^9, 10^. Due to its critical involvement in the initiation of virus infection, the S protein is the major target of neutralizing antibodies (nAbs) and the antigen of choice in vaccine development ^11^. Of the currently approved or authorized SARS-CoV-2 vaccines, four employ two proline substitutions in the S2 domain to prevent its refolding. This prefusion-stabilized construct, referred to as S-2P, is the basis for the Pfizer-BioNTech and Moderna mRNA-based vaccines, Janssen/J&J Ad26-based vaccine and Novavax subunit-based vaccine, and were premised on homologous positions in the MERS-CoV spike resulting in higher titers of nAbs when compared to the wild-type spike ^12^. A fifth SARS-CoV-2 vaccine formulation, ChAdOx1-S, comprises a membrane-anchored wild-type spike protein that retains a trimeric prefusion conformation ^13^. Additional efforts to design spike-based vaccines also involve stabilizing the prefusion conformation of the spike ectodomain reviewed in ^14, 15^. One of the most promising antigens, Hexapro, which comprises six prolines , exhibits higher expression levels and resistance to heat and physical stress than S-2P ^16^, and is the antigen of choice in a Newcastle disease Virus (NDV)-vectored COVID-19 vaccine candidate in phase II/III clinical trial (ClinicalTrials.gov: NCT05354024), as well as in other SARS-CoV-2 spike subunit vaccines ^17, 18^.

Most of the current first-generation vaccines utilize the early pandemic spike protein identified from the Wuhan-1 isolate. However, several mutations have accumulated in the spike protein, resulting in the emergence of variants of concern (VOCs) ^19, 20^. Of particular interest is the omicron variant of SARS-CoV-2, which possesses extensive capabilities to escape from the neutralizing immunity elicited by mRNA-based vaccines ^21–23^, and has reignited debate over the need for booster vaccine doses or reformulated vaccines ^24, 25^. While a third or fourth vaccination dose restores the neutralization of omicron and reduces COVID-19 severity in the short term, the currently used booster approach is unsustainable, warranting the development of vaccines that promote more durable immunity ^21–23^.

While the rapid development of the multiple COVID-19 vaccines has been crucial in curbing the ongoing pandemic, there are several limitations. The mRNA vaccines are expensive and hard to transport due to the freezing requirements. Although the adenovirus vector-based vaccines have greater stability than the mRNA vaccine and have no freezing requirements, The Food and Drug Administration (FDA) and the Centers for Disease Control and Prevention (CDC) have restricted the use of Ad26.COV2 in the United States due to unusual but serious adverse effect of thrombotic events with thrombocytopenia ^26^. A similar risk has not been identified with mRNA vaccines, but myocarditis and pericarditis have been reported ^27–29^. Besides, serum neutralizing antibody titers induced by mRNA are short-lived with half-life of approximately 30 days ^30–32^. Hence, the development of other vaccine platforms and strategies that can elicit a longer-lasting immune response with an acceptable safety profile are highly warranted.

The live-attenuated measles virus (MeV) vaccine is a highly attractive vectored vaccine since it has a proven track record of safety and efficacy in humans and is known to elicit long-lasting B- and T-cell responses, with a reported measles-specific antibody half-life of 3,014 years ^33–35^. This durable protective immune response has been attributed to efficient replication and spread of MeV in the lymphoid tissue followed by persistence of MeV RNA once the infectious virus is eliminated ^36, 37^. Consequently, a MeV-vectored vaccine has the potential to elicit long-lasting immune responses against heterologous antigens. Indeed, the live-attenuated MeV vaccine has been engineered as a vectored vaccine against a variety of pathogens ^38, 39^, and a MeV-based vaccine candidate against Chikungunya has shown promising results in a phase II clinical trial ^40^. Several attempts have been made to use MeV-based vaccines for SARS-CoV-2 ^41–43^. These preclinical candidates were based on the membrane-anchored wild-type spike protein ^42^ or the pre-fusion stabilized spike protein (S-2P) ^41^. A third construct used a secreted form of the S-2P with a self-trimerizing “foldon” domain replacing the transmembrane and cytoplasmic domains of the spike ^43^. Notwithstanding, the clinical development of a counterpart MeV-based SARS-CoV-2 vaccine candidate (V592) was recently discontinued due to low seroconversion rates, especially in measles-immune individuals ^44, 45^. Currently, it is not known the role of the spike design and oligomerization state into the magnitude and breath of the elicited immune response.

Here, we aimed to generate a MeV-vectored vaccine capable of driving high SARS-CoV-2 neutralizing antibody responses. To achieve this, we first sought to circumvent blunting of vaccine efficacy by pre-existing anti-measles antibodies ^45^, using our previously developed MeV-based vaccine with extensively modified surface glycoproteins ^46^. Then, we proceeded to explore the immunogenicity of various measles-vector COVID-19 vaccine candidates expressing genetically modified SARS-CoV-2 spike ectodomain constructs. We demonstrate that artificial trimerization of the SARS-CoV-2 spike protein is necessary for the induction of a robust nAb response in type-I interferon deficient, human CD46 transgenic mice (IFNAR^−/−^-CD46Ge). When we scaffolded the trimeric SARS-CoV-2 spike protein onto the homododecameric neutrophil-activating protein (NAP) from Helicobacter pylori (H. pylori), the resulting construct triggered a significantly higher production of nAbs than the unscaffolded trimeric spike. Furthermore, although two doses of a MeV/COVID-19 vaccine candidate encoding a historical Wuhan spike glycoprotein elicited robust production of nAbs against historical SARS-CoV-2 variants, the titers against the omicron lineage were significantly lower. An omicron-matched MeV/COVID-19 booster increased the nAb responses against both omicron and historical variants. Finally, we show that serum antibodies induced in IFNAR^−/−^-CD46Ge by the MeV/SARS-CoV-2 can protect K18-hACE2 mice from COVID-19 after infection with former and current SARS-CoV-2 variants. These results will inform the further clinical development of MeV/COVID-19 vaccine candidates and therapies.

## Results

### Only the full-length SARS-CoV-2 spike elicits pseudo virus neutralizing antibodies in IFNAR*^−/−^*-CD46Ge mice

Type-I interferon deficient, human CD46 transgenic mice (IFNAR^−/−^-CD46Ge) are considered the gold-standard small animal model for the analysis of rMeV-based vaccine candidates ^38^. Since different mouse strains might differ in their responsiveness to antigen stimuli ^47^, we initially sought to evaluate in this animal model the antigenic properties of the SARS-CoV-2 full-length spike protein and three different subunits vaccines; thus, we analyzed (1) the full-length spike ectodomain (Wuhan-Hu-1 isolate, S1+S2, amino acids 16 to 1213), (2) the S1 domain (amino acids 16 to 685), (3) the S2 domain (amino acids 686 to 1213) and (4), the receptor-binding domain (RBD, amino acids 319 to 541). We immunized IFNAR^−/−^-CD46Ge mice twice at 3-week intervals with 5 μg of recombinant protein adjuvanted with aluminum hydroxide gel (Alum) via the intraperitoneal route. Alum was chosen as it is the most used adjuvant in vaccines in humans. Serum samples were then collected on days 21 (before boost) and 42 and analyzed by ELISA for antibodies binding to various spike proteins and domains. After a first immunization, the levels of binding antibodies were low to undetectable, but they were significantly increased after a second dose (Figure 1A). An exception was observed in antisera generated in response to S1-RBD, which exhibited no binding to S1+S2 (Figure 1A). When we analyzed these sera in more detail, we found that both the full-length spike ectodomain (S1+S2) and the S2 subunit elicited the production of IgG antibodies with comparably strong binding to both S1+S2 and S2 alone (Figure 1B). In contrast, both the S1-RBD and the S1 domains induced the generation of antibodies that were specific to the S1-RBD (Figure 1B). These results indicate that the RBD is immunodominant in the S1 domain, although most epitopes are located within the S2 subunit. Alternatively, the lower immunogenicity of the S1 and S1-RBD could be related to the loss of structural epitopes in the truncated soluble forms.

**Figure 1.**
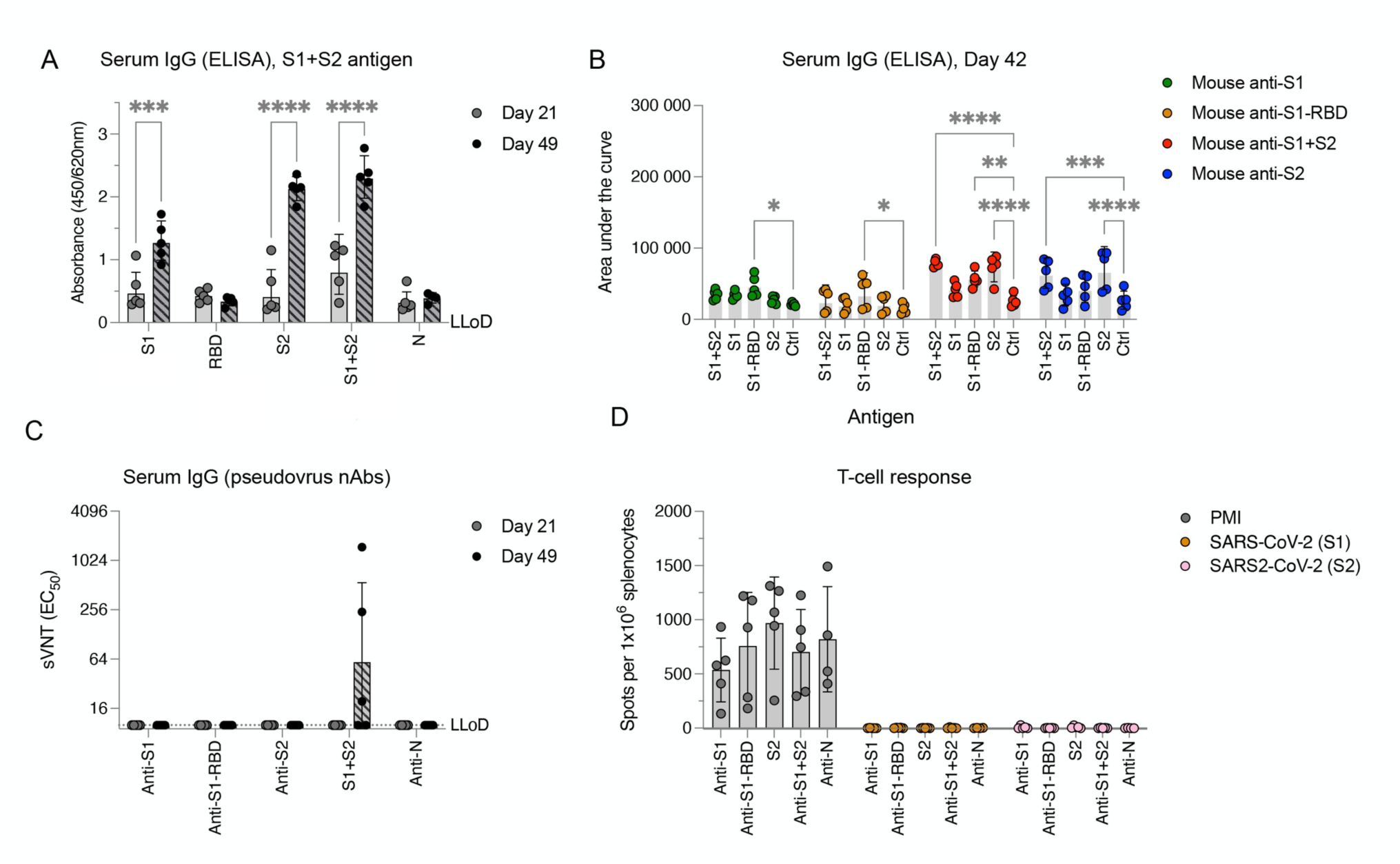
Full-length SARS-CoV-2 spike ectodomain protein elicits poor pseudovirus-neutralizing antibody production and does not elicit T-cell responses. (A) SARS-CoV-2-spike binding responses. IFNAR^−/−^-CD46Ge mice were vaccinated intraperitoneally at days 0 and 21 with 5 µg of purified SARS-CoV-2 protein and alum adjuvant: full-length spike ectodomain (S1+S2), spike receptor-binding domain (S1-RBD), Spike S1 domain (S1), spike S2 domain (S2), and nucleocapsid (N). Serum samples were collected on days 21 (before the second vaccination) and 41 for quantification of the levels of Spike ectodomain binding IgG antibodies by enzyme-linked immunosorbent assay (ELISA). (B) Binding IgG in serum from mice vaccinated twice with SARS-CoV-2 spike protein or domains (S1, green; S1-RBD, orange; S1+S1, red; S2, blue) was also quantified by ELISA for binding to homologous or heterologous antigens. Serial fivefold dilutions were assessed, and data were computed as the area under the curve. (C) Pseudovirus-neutralizing antibody responses. Neutralizing-antibody titers in mice vaccinated once (day 21) or twice (day 42) with the indicated SARS-CoV-2 proteins were determined using pseudotyped viruses expressing the SARS-CoV-2 spike protein bearing the D614G amino acid change. Virus neutralization was plotted as the percentage of relative virus infection over the inverse of serum dilution. The inverse of the serum dilution resulting in 50% inhibition of infection (EC50) was determined and plotted. Antibody titers below the lower limit of detection (LLoD) were treated as 0.5xLLoD. (D) T-cell responses elicited against SARS-CoV-2 spike. An ELISPOT assay for IFN-γ was performed on splenocytes isolated from mice vaccinated twice (day 42) and stimulated ex vivo with PMA/ionomycin (PMI) or two separate pools of 15-mer, 11-aa-overlapping peptides comprising the SARS-CoV-2 spike (S1, aa 1-632; S2, aa 632-1273). The data are shown as IFN-γ-secreting cells or spot-forming cells (SFCs) per 1x10^6^ splenocytes. Values represent the geometric mean ± geometric standard deviation, with each data point representing an individual mouse. Statistical significance was determined using two-way ANOVA with Dunnett’s multiple comparison test (*, p<0.05; **, p<0.005; ***, p=0.0005; ****, p<0.0001).

nAb responses against SARS-CoV-2 were next measured using a previously described lentiviral pseudotype assay ^48^. Neutralization activity was only observed in antisera generated in response to the full-length S1-S2 ectodomain. However, the nAb titers were low and were detected in only three out of five animals (Figure 1C). Thus, antibodies produced in response to the soluble and purified full-length spike protein target recognized predominantly nonneutralizing epitopes.

Finally, we assessed the ability of the various spike domains to induce a T-cell response. To this end, splenocytes from immunized animals were collected three weeks after the booster was administered, and the cells were analyzed by ELISPOT assay for antigen-specific IFN-γ production. While a similar basic reactivity to nonspecific T-cell stimulation was observed across different groups, no reactivity was observed when splenocytes were stimulated ex vivo with two different pools of SARS-CoV-2 spike peptides (Figure 1D). Taken together, these data suggest that the full-length SARS-CoV-2 spike protein was the only antigen able to elicit an immune response, and that it exclusively engaged the humoral arm of the immune response and presented narrow neutralizing activity.

### Multimerization of the SARS-CoV-2 spike protein enhances the pseudovirus-neutralizing antibody response

A desirable property of a human vaccine is the ability to induce nAbs. Research in other type I fusion glycoproteins has suggested that nAbs recognize metastable quaternary epitopes rather than monomer forms ^49–51^. Therefore, we postulated that the high ratio of binding antibodies to nAbs that was observed when the soluble purified protein was used for immunization, may have been related to the lack of a quaternary assembly of the prefusion trimer. Also suggesting that nAbs recognize a quaternary spike epitope in the metastable prefusion conformation, as observed in other type I fusion glycoproteins ^12, 52–54^. To begin to test our hypothesis, we incorporated a self-trimerizing T4 fibrin motif (foldON) ^55^ in the full-length spike ectodomain in conjugation with a mutated furin cleavage site in the spike and the previously reported stabilizing six proline substitutions in the spike (HexaPro, S-6p) ^56^, which disfavor formation of an extended coiled coil ^14^. Additionally, we produced a genetic fusion at the C-terminus of the SARS-CoV-2 spike protein with H. pylori NAP (Figure 2A). NAP is a 27-nm-wide dodecameric protein with four 3-fold axes ^57, 58^, a feature that could enable multivalent display of immunogens on the exterior surface ^59–63^. Next, both trimeric and full-length spike ectodomain (S1+S2, herein termed SARS-CoV-2S6p3) and SARS-CoV-2Sp3-NAP (herein termed SARS-CoV-2S6p312) were recombinantly expressed using mammalian cells to ensure the proper folding and glycosylation pattern of the proteins. SDS‒PAGE analysis followed by Coomassie blue staining of purified SARS-CoV-2S6p3 and SARS-CoV-2S6p312 revealed apparent molecular weights of 180 kDa and 210 kDa, respectively, under reducing conditions, which suggested proper genetic fusion of NAP (Supplementary Figure 1A). This analysis also revealed that the preparations were of high purity. Further native gel electrophoresis demonstrated that both SARS-CoV-2 spike constructs preferentially assembled as mature trimers, as expected from a correctly fused foldON domain (three ∼270 kDa units, Supplementary Figure 1B). The purity and homogeneity of the recombinant proteins was also verified by negative transmission electron microscopy (negative-TEM). We found after NAP conjugation a higher degree of protein aggregation, consistent with nanoparticle display (Supplementary Figure 2).

**Figure 2.**
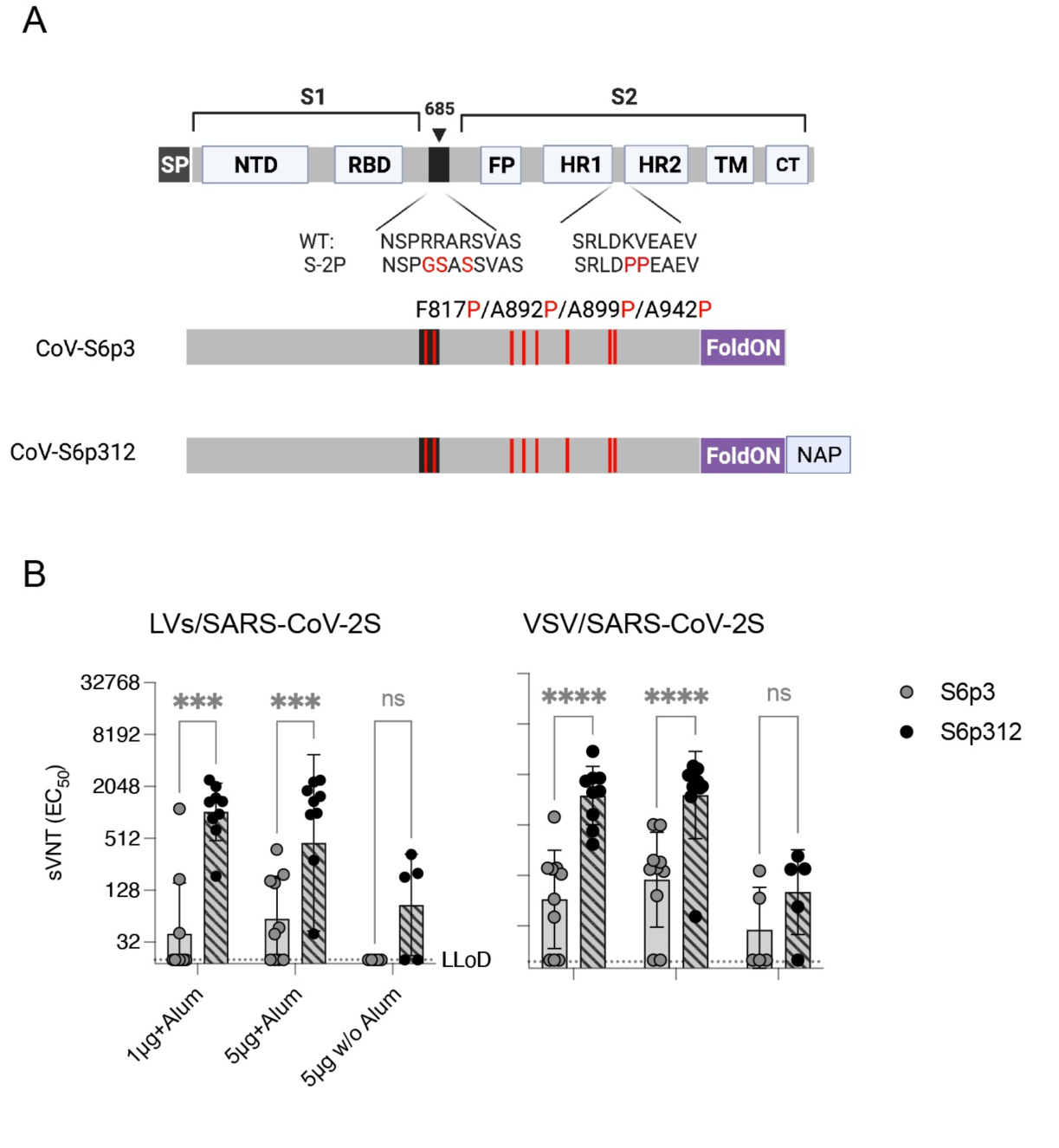
Multimerization of SARS-CoV-2 spike enhances neutralizing antibody responses. **(A)** Schematic diagram of the full-length SARS-CoV-2 spike and engineered full-length ectodomain spikes. Some of the structural domains shown include the cleavable signal peptide (SP), N-terminal domain (NTD), receptor-binding domain (RBD), S2 cleavage site (685, black), fusion peptide (FP), heptad repeats 1 and 2 (HR1 and HR2), transmembrane domain (TM) and cytoplasmic tail (CT). The native furin cleavage site was altered (RRAR→GSAS) to inhibit proteolytic cleavage, and six prolines, noted in red text, were introduced to further increase stability. Further modifications include the C-terminal domain of the T4 fibritin (foldON, purple) placed at the C-terminus of the spike and the H. pylori neutrophil-activating protein (NAP, blue) preceded by a GlySer linker. (B) Pseudovirus-neutralizing antibody responses. Mice were vaccinated once with either 1 or 5 μg of alum-adjuvanted proteins, and neutralizing antibodies in serum samples collected 21 days post-vaccination were quantified using LV-SARS-CoV-2 pseudoviruses (left panel) and VSV-SARS-CoV-2-S pseudoviruses (right panel). Antibody titers below the lower limit of detection (LLoD) were replaced with 0.5xLLoD. Black dots represent individual mice, and bars and error bars depict the geometric mean ± geometric standard deviation, respectively. Statistical analysis among groups was calculated by two-way ANOVA with Bonferroni’s post test (ns, p>0.05; ****, p<0.0001).

Finally, we compared the immunogenicity of these spike proteins by vaccinating 5-10 IFNAR^−/−^-CD46Ge mice with alum-adjuvanted formulations containing 1 μg or 5 μg of SARS-CoV-2 spike protein or with 5 μg of SARS-CoV-2 spike protein without alum. Serum samples were then collected at week 3 to measure the levels of pseudovirus nAbs. Mirroring the data presented above, not all the animals vaccinated with a prefusion, trimeric SARS-CoV-2 (SARS-CoV-2S6p3) produced neutralizing antibodies despite the use of the adjuvant (3/10 for the 1 μg dose and 6/10 for the 5 μg dose), and those that did, exhibited low geometric mean titers (GMTs), i.e., 199 and 123. In contrast, all animals vaccinated with a homologous but covalently linked NAP-tagged SARS-CoV-2 spike (SARS-CoV-2S6p312) exhibited seroconversion when alum was used as an adjuvant and GMTs that were roughly 5-fold higher, i.e., 1,038 and 453 for the 1 μg and 5 μg dose, respectively (Figure 2B, left panel). We observed comparable results when neutralization titers were measured using VSV-SARS-CoV-2-S pseudoviruses (Figure 2B, right panel). These data strongly suggest that the conformation of relevant B-cell epitopes is likely to be preserved in the metastable prefusion and stable postfusion products. We conclude from this experiment that recombinant SARS-CoV-2 spike is poorly immunogenic, but its multivalent display on a self-assembling nanoparticle scaffold markedly improves its immunogenicity.

### Measles Virus-based SARS-CoV-2 candidates express spike proteins at comparable levels

As only the full-length spike ectodomain was able to elicit the production of nAbs in IFNAR^−/−^-CD46Ge mice, we next sought to generate rMeV-based SARS-CoV-2 vaccine candidates expressing various full-length SARS-CoV-2 spike proteins. We have recently reported the generation of a remodeled Moraten-based MeV (MeV-MR) with reduced susceptibility to neutralization by anti-MeV antibodies ^46^. Since most individuals are seropositive for measles, which has impacted the immunogenicity of a previously developed measles-vectored SARS-CoV-2 vaccine candidate ^45^, we selected MeV-MR as our vector platform. A panel of rMeV-MR constructs encoding unmodified or modified versions of the spike was cloned between the MeV-P- and MeV-M-coding sequences of MeV-MR ^46^. Among the modifications of the spike that were generated, we replaced the native signal sequence with the murine IgG **κ** leader sequence followed by a hemagglutinin (HA) tag. This signal peptide has been shown to significantly enhance the immunogenicity of an adenoviral vectored vaccine platform ^64^. Additionally, we included two sets of prefusion-stabilized forms of spike, the S-2P construct ^65^ and the S-6P ^16^, for comparison. At the time this work was being conducted, it was unknown which form of pre-fusion spike was more immunogenic. Finally, we included the product of genetic fusion of NAP with or without the presence of the foldON domain.

Altogether, we designed seven different constructs (Figure 3A): (i) wild-type leader sequence with deletion of the S cytoplasmic tail (CoV-SΔCT); (ii) the CoV-SΔCT protein with alteration of the furin cleavage site and six stabilizing proline substitutions (CoV-S6ΔCT); (iii) the CoV-ΔCT protein with an IgGκ leader sequence and deletion of the spike transmembrane region, reflecting the soluble ectodomain, fused to NAP (CoV-S-12); (iv) The CoV-S-12 protein with alteration of the furin cleavage site, two stabilizing proline substitutions and a STOP codon before NAP (CoV-S2p); (v) the CoV-S2p protein without a STOP codon before NAP (CoV-S2p12); (vi) the CoV-S2p12 protein with a foldON trimerization motif between the pre-fusion spike and NAP (CoV-S2p312); and (vii) the CoV-S2p312 protein with four additional proline substitutions (CoV-S6p312).

**Figure 3.**
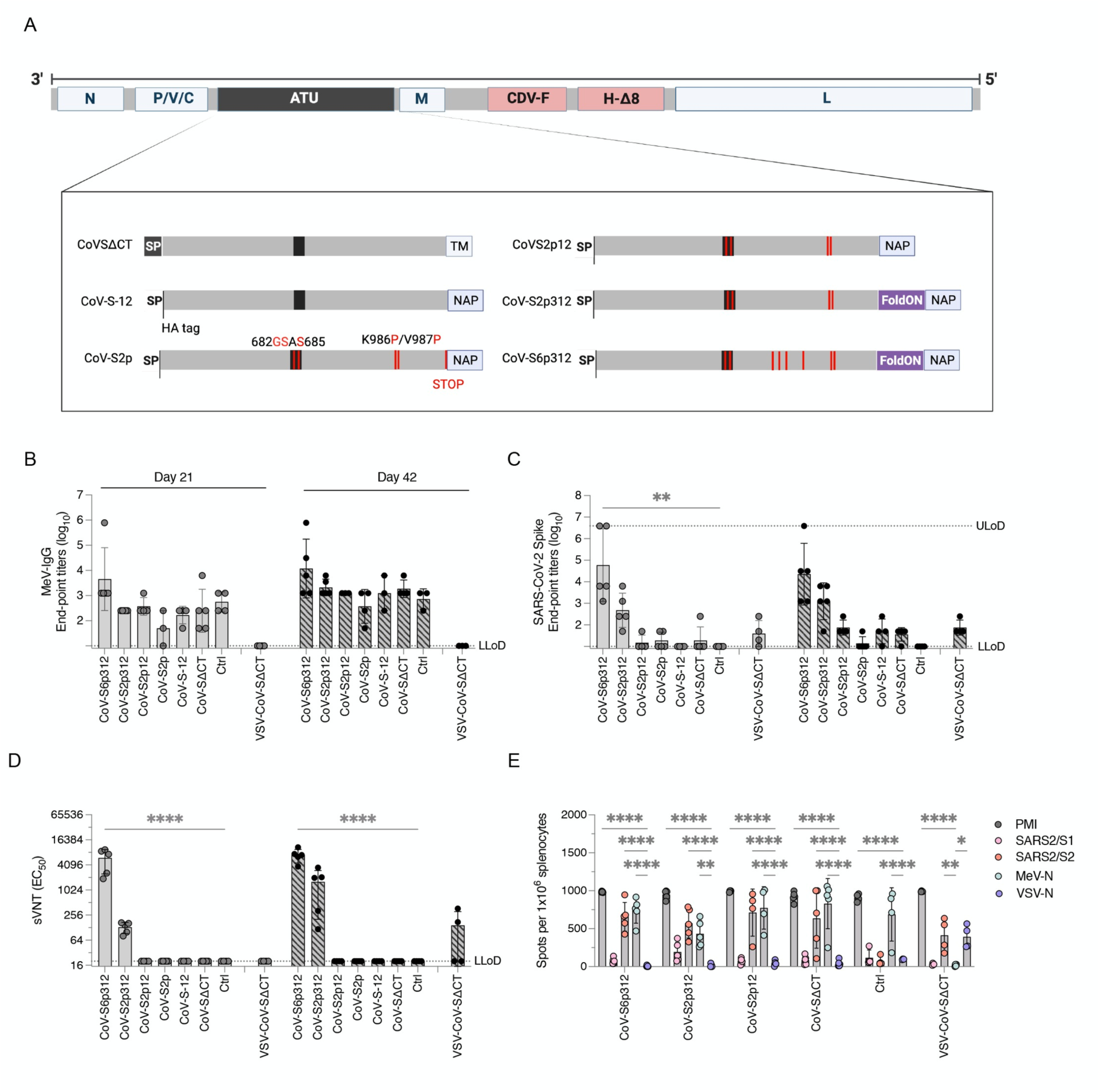
Trimerization and stabilization of SARS-CoV-2 spike constructs augment the humoral antibody response. **(A)** Schematics of the MeV-MR vector and SARS-CoV-2 spike-based constructs inserted in it as an additional transcript unit (ATU), labeled as in Figure 2. The MeV genome consist of the following genes from the Moraten vaccine strain: nucleoprotein, phosphoprotein, V and C accessory proteins, matrix, and large polymerase protein. The envelope glycoproteins were substituted for canine distemper virus fusion protein and a wild-type hemagglutinin protein with deletion of 8 antigenic sites. The schematics show below depict modifications to the SARS-CoV-2 spike protein, including deletions of the transmembrane and/or cytoplasmic tail region as well as the substitution of the SARS-CoV-2 spike signal peptide by the murine IgG kappa leader sequence, followed by an HA tag. Among other modifications, Helicobacter pylori NAP was genetically fused at the extreme C-terminus of the spike and either preceded or not by a stop termination codon. Alternatively, a foldON trimerization domain was inserted between the spike and NAP. **(B and C)** Binding IgG responses. IFNAR^−/−^-CD46Ge mice were vaccinated intraperitoneally at days 0 and 21 with 1x10^5^ pfu of either rMeV or vesicular stomatitis virus (VSV) expressing various spike-based constructs. Serum samples were collected on days 21 (before the second vaccination) and 42 and assessed by enzyme-linked immunosorbent assay (ELISA) for IgG binding to **(B)** MeV-bulk antigen and **(C)** the spike ectodomain. **(D)** Pseudovirus-neutralizing antibody responses. Neutralizing antibody titers in mice vaccinated once (day 21) or twice (day 42) with the indicated recombinant virus were determined using pseudotyped lentiviruses expressing the SARS-CoV-2 spike D614G construct, as previously shown in Figure 1. Virus neutralization was plotted as the percentage of inhibition of virus infection relative to that of virus incubated with negative mouse serum over the inverse of the serum dilution. The inverse of the serum dilution resulting in 50% inhibition of infection (EC50) is plotted. **(E)** ELISPOT assay for IFN-γ by splenocytes isolated from mice vaccinated twice (day 42) and stimulated ex vivo with PMA/ionomycin or antigen-specific peptides. The number of spot-forming cells (SFC) per 1x10^6^ splenocytes is plotted. Values represent the geometric mean ± geometric standard deviation, with each data point representing an individual mouse. Statistical significance was determined using two-way ANOVA with Dunnett’s multiple comparison test (*, p<0.05; **, p<0.01; ****, p<0.0001).

All rMeVs were rescued and propagated in Vero cells to produce virus stocks, each reaching comparable titers (∼ 10^6^ pfu/mL). Next, virus integrity was assessed by full-genome NGS. Whereas the MeV coding sequences were identical, some amino acid changes were noted in the spike region of some of the rMeVs. In the CoV-S6ΔCT construct 15 nonengineered amino acid changes in addition to an early stop termination were detected due to a single point mutation (Supplementary Figure 3). Also, a single A890V amino acid substitution was present in the CoV-S6p312 virus. No amino acid changes were observed in any of the other viruses. These results suggest that the nonfusogenic versions of the spike protein are subjected to selection pressure when displayed on the MeV coat. Consequently, the rMeV-MR-CoV-S6ΔCT vaccine candidate was abandoned, and no further characterization was performed.

Finally, the expression of the spike protein was analyzed by western blot analysis of Vero cells infected with rMeVs (Supplementary Figure 4). Similar MeV-N antigenic material was detected for all the rMeVs, suggesting similar kinetic growth among the viruses. When the cells were infected with rMeV expressing SARS-CoV-2 spike with prefusion-stabilizing amino acid changes (S2p or S6p), the full-length spike proteins were detected with antibodies against the SARS-CoV-2 spike and against the N-terminus HA-tag. Among these constructs, and in the absence of C-terminal NAP, the recombinant spike protein was mostly expressed in a soluble form and was secreted into the culture medium (MR-CoV-S2p). When NAP was genetically added to the C-terminus of the spike, the spike was detected in both the culture medium and the cell pellet, and an additional band >250 kDa was detected (CoV-S2p, CoV-S2p312, CoV-S6p312). In cells infected with MR-CoV-SΔCT, both full-length and cleaved spike proteins were detected exclusively with the SARS-CoV-2 spike antibody, and they were detected predominantly in the cell pellet. Unlike the soluble and prefusion-stabilized constructs, for which low band intensity was noted when an anti-SARS-Co-V-2 spike antibody was used, a prominent signal was observed for the SΔCT construct. Overall, these results indicate that rMeV can express various recombinant spike proteins and that both oligomerization status and epitope accessibility vary among the constructs.

### Artificial trimerization of the spike protein is critical for the immunogenicity of MeV-based SARS-CoV-2 vaccine candidates

We next evaluated the immunogenicity of the different vaccine candidates. To this end, 1x10^5^ plaque-forming units of the various viruses were used to vaccinate 8- to 12-week-old IFNAR^−/−^-CD46Ge mice on days 0 and 21. Serum samples were then collected on days 21 (before boost) and 42 to assess the presence of S- and MeV-specific IgG antibodies by ELISA. As a negative control for vaccination, we used an isogenic MeV-MR encoding an irrelevant antigen or a VSV-G protein-pseudotyped VSV expressing SARS-CoV-2 spike [VSV-CoV-SΔCT]^66^.

End-point titers of sera isolated from animals vaccinated with rMeV constructs, including the control, exhibited antibodies that bound to MeV antigens that were detectable 3 weeks after the first vaccination. The levels of these antibodies increased by more than one log after a second dose with the homologous virus, indicating the occurrence of vaccine-induced responses in all animals (Figure 3B). Even though IgG antibodies specific to MeV were detected in animals that received MeV-MR, seroconversion to SARS-CoV-2 was not observed in all groups even after two doses. Specific IgG antibodies to SARS-CoV-2 spike were detected in 100% of animals vaccinated once with rMeV expressing a trimeric and stabilized SARS-CoV-2 spike, CoV-S2p312 or CoV-S6p312 and in 100% of animals vaccinated twice with rMeV-MR-CoV-S2p12 and VSV-CoV-SΔCT (Figure 3C).

The neutralizing activity of the antibodies was measured using SARS-CoV-2 spike-pseudotyped lentiviruses. nAbs were only detected in mice vaccinated with trimeric and stabilized SARS-CoV-2 spike (CoV-S2p312 and CoV-S6p312) and not in mice vaccinated with any of the other rMeVs. In the CoV-S2p312 and CoV-S6p312-vaccinated mice, nAbs were detected after one dose, whereas two doses of VSV-CoV-SΔCT were required for neutralizing activity to reach detectable levels in some of the vaccinated animals (Figure 3D). Among animals vaccinated with trimeric and stabilized SARS-CoV-2 constructs, the largest difference was observed in their pseudovirus-neutralizing activity, for which one dose of MR-CoV-S6p312 was superior to two doses of MR-CoV-S2p312 (Figure 3D).

Cell-mediated immunity was additionally assessed on day 42 by ELISPOT analysis. All animals that had been vaccinated with any rMeV showed a strong IFN-γ-producing T-cell response (Figure 3E). Similarly, the splenocytes isolated from all animals that received viral vectors expressing SARS-CoV-2 constructs showed reactivity to SARS-CoV-2 peptides, even in animals that previously failed to mount a SARS-CoV-2-specific IgG response. When two pools of SARS-CoV-2 spike were used to stimulate the splenocytes, we observed a higher level of IFN-γ-producing T cells specific for peptides spanning the S2 subunit (aa 633 to 1258). We conclude from these results that the SARS-CoV-2 spike protein has an intrinsic propensity to hamper B-cell responses and that trimerization is key for the induction of anti-SARS-CoV-2 spike IgG antibody production, where prefusion-stabilizing mutations augment the induction of neutralizing responses.

### Immunity elicited by MeV/SARS-CoV-2 is Th1 polarized

Vaccine-associated enhanced respiratory pathology after SARS-CoV-2 infection correlates with a Th2-biased immune response ^67, 68^. Since profiling of IgG1 and IgG2a isotypes can serve as an indication of T-cell polarization ^69^, we next measured the levels of two IgG subclasses of SARS-CoV-2 spike-specific antibodies by ELISA. As a control for a Th2-skewed response, we used serum from mice immunized twice with alum-adjuvanted SARS-CoV-2 spike protein ^42^. In these mice, a significant (p<0.05) difference in the levels of IgG1 and IgG2a subclasses was observed, with IgG1 levels being significantly higher than IgG2a levels (Figure 4A, right panel). In contrast, mice vaccinated with MeV-MR-CoV-S6p312 produced comparable antibody IgG1 and IgG2a titers after one dose. After two doses, a statistically significant predominance of IgG2a was observed, indicative of a Th1-skewed response (Figure 4, left panel). We further confirmed the Th1/Th2 balance by analyzing the cytokine profile of splenocytes stimulated with a SARS-CoV-2 spike peptide pool. Specifically, splenocytes obtained from vaccinated animals were treated with DMSO or the SARS-CoV-2 spike peptide pool, and cytokine secretion was quantified in the cell culture supernatant with a ProcartaPlex multiplex panel. The results showed strong Th1 polarization, based on the production of IL-1β, IL-2, IL-12, TNF-α, and IFN-γ (Figure 4B). No Th2-associated cytokines (IL-5, IL-4 and IL-13) were detected. Collectively, the assessments of both humoral and cellular responses revealed a desirable Th1-biased immune response elicited by MR-CoV-S6p312 .

**Figure 4.**
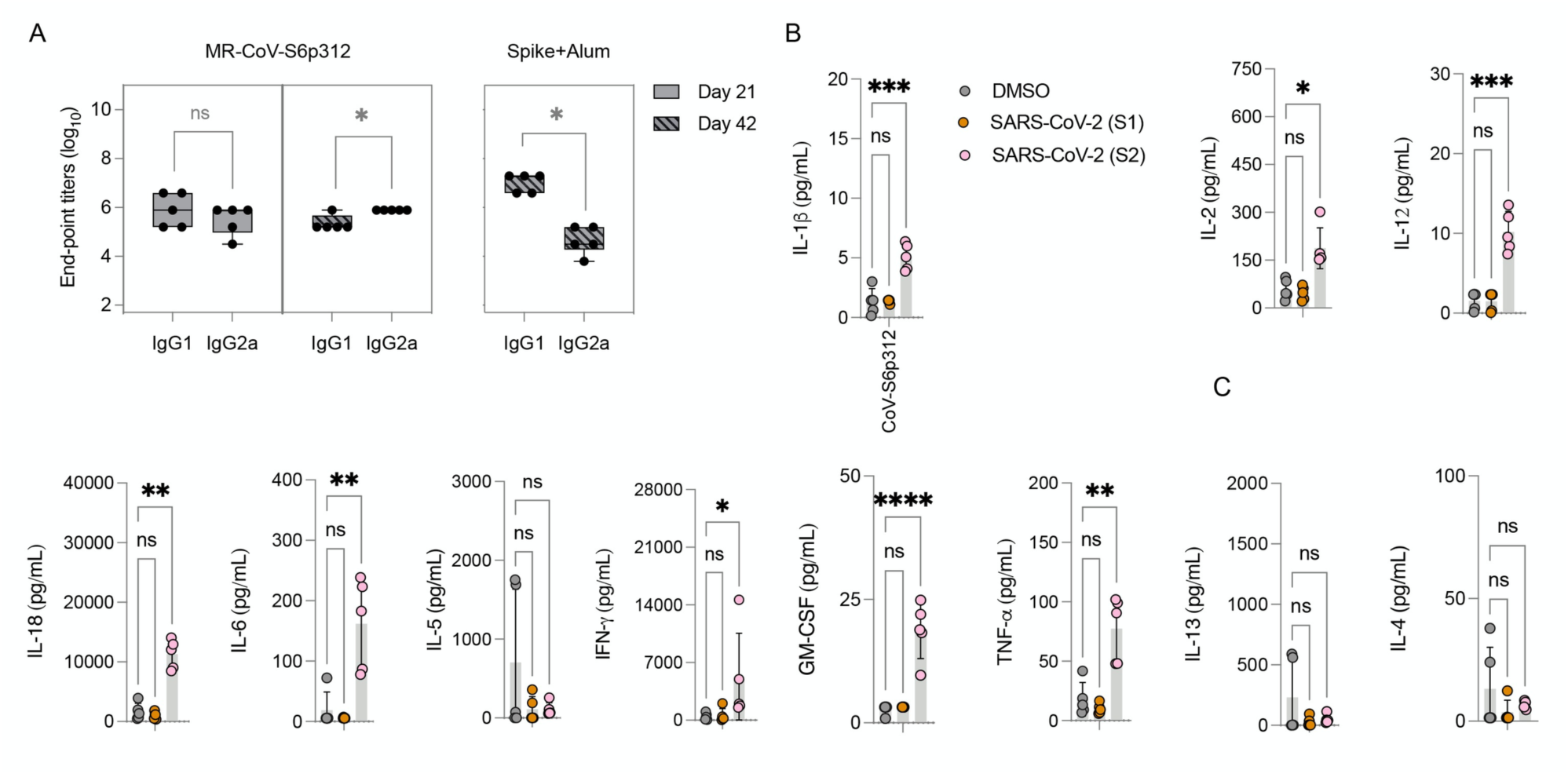
MR-CoV-S6p312 elicits a Th1-oriented humoral and cellular immune response. (A) Isotype analysis of anti-SARS-CoV-2 spike antibodies. Serum samples from IFNAR^−/−^-CD46Ge mice vaccinated once (day 21) or twice (day 42) with MR-CoV-S6p312 were analyzed by ELISA for IgG1 and IgG2a antibody binding to SARS-CoV-2. Serum from mice vaccinated twice with purified SARS-CoV-2 Spike adjuvanted with alum was used as a control for the Th2-biased humoral response. **(B)** Cytokine production from the splenocytes of vaccinated mice. Splenocytes isolated from vaccinated mice were stimulated as indicated in Figure 1, and cytokine secretion in the supernatant was analyzed by multiplex cytokine analysis. Dots represent individual animals, and horizontal bars and error bars are the mean ± SD. IL-1b lower limit of detection (LLOD): 1.45 pg/mL; IL-12 LLoD: 1.68 pg/mL; TNF-a LLoD 3.48 pg/mL; IFN-γ LLoD 2.19 pg/mL; GM-CSF LLoD: 3.20 pg/mL;IL-6 LLoD: 5.52 pg/mL; IL-5 LLoD: 2.19 pg/mL; IL-2 LLoD 1.88 pg/mL;IL-4 LLoD: 1.37 pg/mL; IL-13 LLoD: 2.86 pg/mL. Statistical significance was determined using two-way ANOVA with Dunnett’s multiple comparison test (*, p<0.05; **, p<0.003; ***, p<0.0003; ****, p<0.0001).

### MeV/SARS-CoV-2 vaccine based on the historical spike protein elicited low neutralizing antibodies against some SARS-CoV-2 variants

We next investigated whether this favorable Th1-type immune response elicited in MeV-vaccinated animals could neutralize SARS-CoV-2 variants, which have almost completely replaced the original SARS-CoV-2 that was used for our vaccine design. As of January 2022, the World Health Organization had defined five VOCs (alpha [B.1.17], beta [B1.351], gamma [P1], delta (B.1.617.2], omicron [B.1.1.529]) as well as five variants of interest (iota [B.1.526], kappa [B.1.617.1], lambda [C.37], mu [B.1.621], epsilon [B.1.427/B.1.429]). Among the variants of greatest concern is the omicron variant (formerly known as B1.1.529), which exhibits 34 amino acid substitutions in the spike protein, in contrast with the 8-12 modifications that were observed in previous VOCs. This finding has resulted in partial or complete escape from the humoral ^70, 71^ (Supplementary Figure 5) but not T-cell responses elicited upon vaccination with different platforms ^70–75^. Our primary approach was to perform antibody neutralization assays with pseudoviruses expressing the omicron BA.1 variant SARS-CoV spike containing the lineage-defining amino acid changes A67V, deletion (Δ)H69-V70, T95I, G142D, ΔV143-Y145, ΔN211, L212I, ins214EPE, T547K, D614G, H655Y, N679K, P681H, G339D, S371L, S373P, S375F, K417N, N440K, G446S, S477N, T478K, E484A, Q493R, Q498R, N501Y, Y505H, N764K, D796Y, N856K, Q954H, N969K, and L981F. Moreover, we generated additional pseudoviruses harboring the spike proteins of other variants. Sera from mice vaccinated with a single vaccine dose of MR-CoV-S6p312 had similar neutralizing effects (p>0.05) on pseudoviruses harboring spikes from alpha, kappa, epsilon and beta. However, a partial or complete loss of neutralization was observed for pseudoviruses harboring spikes from delta, gamma, lambda, omicron BA.1, mu and iota (Figure 5). Hence, some variants are resistant to neutralization by antibodies produced in response to a single dose of our measles-vectored COVID-19 vaccine candidate.

**Figure 5.**
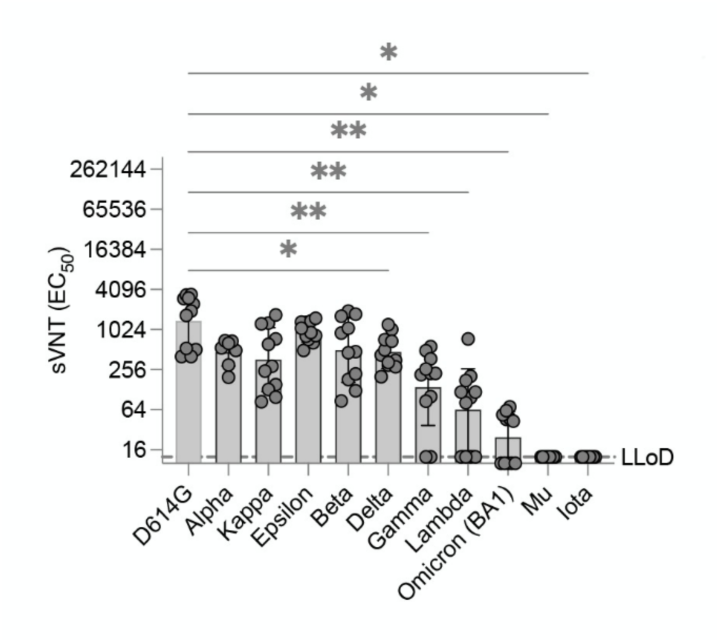
Antibodies elicited by MR-CoV-S6p312 are sensitive to amino acid substitutions present in SARS-CoV-2 variants. Neutralizing activity against SARS-CoV-2 variants. Serum samples from animals vaccinated once with MR-CoV-S6p312 were assessed for neutralizing antibody responses against pseudoviruses bearing the SARS-CoV-2 spike from different variants. Black dots represent individual mouse sera, and bars and error bars depict the geometric mean ± geometric standard deviation, respectively. Statistical analysis among groups was calculated by one-way ANOVA with Dunnett’s post test (ns, p>0.05; *, p<0.05; ****, p<0.0001).

### An Omicron-matched MeV/SARS-CoV-2 vaccine candidate restores neutralizing antibody titers against historical and BA.1 variants

Because the neutralizing antibody response against omicron lineage variant BA.1 was low or absent after a single vaccination with MR-CoV-S6p312 , we sought to evaluate whether (i) there was a constriction of immunity elicited by the original (wt) MR-CoV-S6p312 vaccine candidate and (ii) a homologous wt MR-CoV-S6p312 or a heterologous omicron BA.1-matched MR-CoV-S6p312 vaccine candidate could broaden neutralizing responses against the omicron variant, and if there was a difference between homologous and heterologous boosting.

To address these questions, we sequentially vaccinated two cohorts of 15- to 17-week-old IFNAR^−/−^-CD46Ge mice at weeks 0 and 10 with two doses of wt or one dose of wt and another dose of the BA.1-matched MR-CoV-S6p312 vaccine candidate (Figure 6A). We used a 10-week period between vaccination doses to ensure the presence of affinity-mature, class-switched memory B cells and long-lived plasma cells. We collected blood samples and measured VSV-SARS-CoV-2-S pseudovirus nAbs at peak levels (week 3) ^76^, before boosting (week 10), and 3 weeks thereafter (3 weeks post-boosting). There was significantly less Wuhan strain-neutralization activity measured in the 10-week serum samples than in the 3-week serum samples (3-fold, p<0.0005) in wt construct-vaccinated mice (Figure 6B). Confirming our previous results, the levels of omicron nAbs were low or absent in these animals (Figure 6C). A homologous wt booster shot significantly augmented the levels of Wuhan strain nAbs (p<0.0005), reaching levels comparable to those in week 3 after the first dose (Figure 6B, left panel). Although omicron strain nAbs were detected in all the animals at this point, the GMT of neutralization activity was low, i.e., (1/dilution) ± SEM of (72.8 ± 11.0) (Figure 6C, left panel). However, an omicron-based booster shot not only augmented antibody titers against the omicron variant but also rescued antibody titers against Wuhan strain pseudoviruses (Figure 6B and 6C, right panels). The neutralization titers for Wuhan strain pseudovirus were equivalent (p>0.05) between animals receiving wt or omicron-based boosters. Together, these data strongly suggest that omicron could be considered a SARS-CoV-2 serotype ^77^ and hence that an omicron-based booster is better suited to restore neutralizing antibody titers against not only the homotypic virus but also historical SARS-CoV-2 variants.

**Figure 6.**
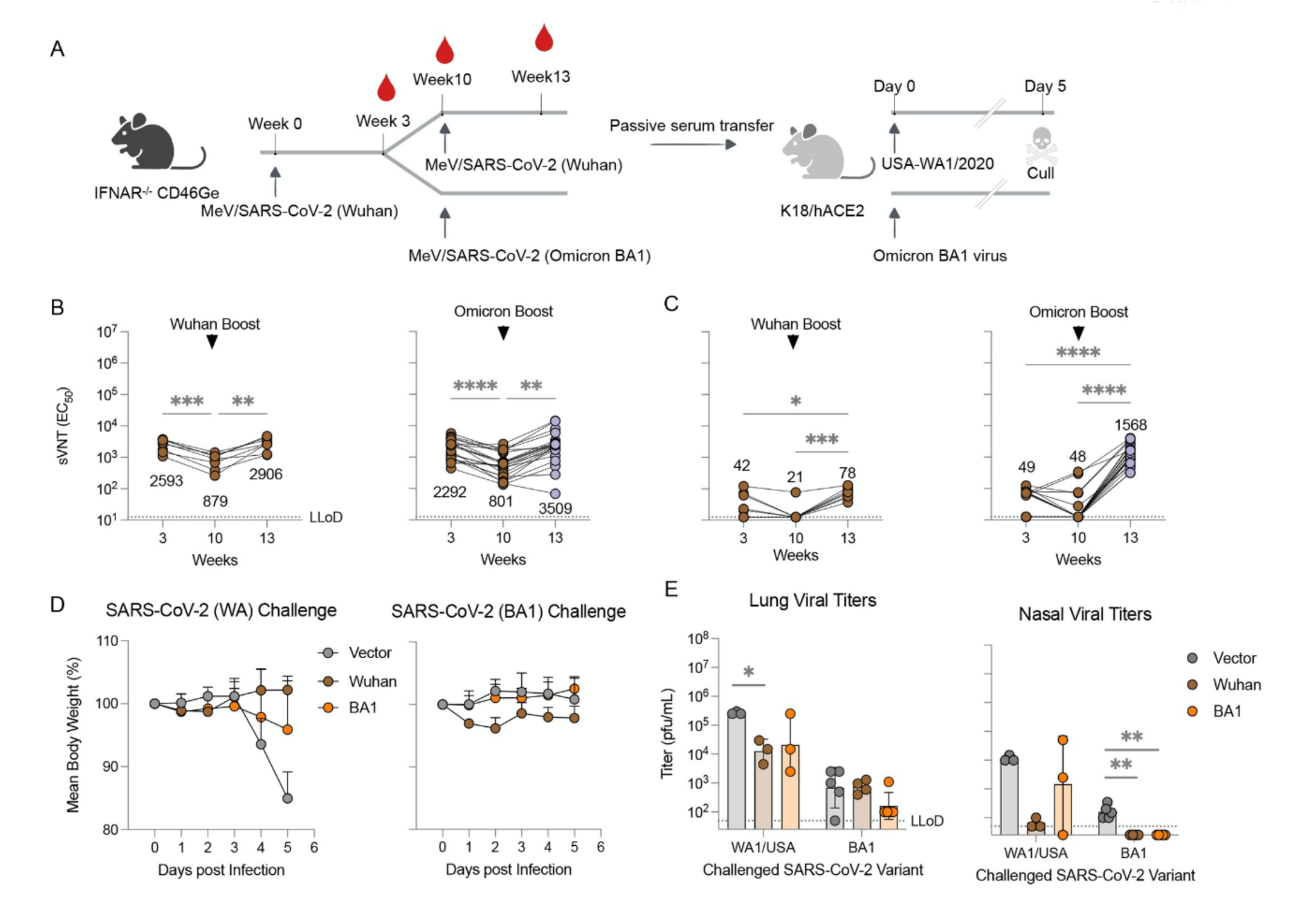
A booster dose of an omicron BA.1-matched MeV/COVID-19 vaccine candidate enhances neutralizing activity and confers protection in K18-hACE2 mice. (**A-C**) Experimental design and serum neutralizing antibodies. (A) IFNAR^−/−^-CD46Ge mice were vaccinated on week 0 with D614G-based MeV/SARS-CoV-S6p312 and boosted on week 10 with the same D614G-based vaccine or an omicron BA.1-based MeV/SARS-CoV-S6p312 vaccine. Serum samples were collected at weeks 3, 10 and 13 and analyzed for pseudovirus-neutralizing antibodies using a Wuhan strain-based VSV/SARS-CoV-2 pseudovirus based on the Wuhan spike (**B**) or omicron BA-1 spike (**C**). Each dot represents an individual mouse sera. Statistical analysis among time-points was calculated by one-way ANOVA with Turkey’s post test (*, p<0.05; **, p<0.01***, p<0.001). **(D)** Protection from body weight loss in mice challenged with SARS-CoV-2 (WA1/2020). K18-ACE2 mice were passively immunized intraperitoneally with serum samples from the previous IFNAR^−/−^-CD46Ge vaccinated animals after a homologous (Wuhan) or heterologous (omicron) boost. Serum samples from animals vaccinated twice with an empty MeV-MR vector were used for sham vaccination (Empty). Two hours later, K18-ACE2 mice were challenged intranasally with WA1/2020 (Wuhan-like strain) or hCoV-19/USA/NY-MSHSPSP-PV44476/2021 (a BA.1 strain) and monitored for body weight loss. **(E)** Virus burden in homogenates from lung and nasal turbinates at 5 days post-challenge with WA1/2020 or Omicron BA.1 virus was assessed by plaque assay. Virus titers below the lower limit of detection (LLoD) were replaced with 0.5xLLoD. Dots represent individual animals, and horizontal bars and error bars are the geometric mean ± geometric standard deviation. Statistical significance between groups was calculated by one-way ANOVA with Dunnett’s’s post test (*, p<0.05; **. p<0.01).

### Vaccine-elicited antibodies from historical and Omicron-matched MeV/SARS-CoV-2 vaccine candidates protect against SARS-Cov-2 challenge

As the in vitro neutralizing activity of our antibodies was promising, we next tested whether this extended to similarly strong in vivo activity against SARS-CoV-2. To determine the protective efficacy of homologous and heterologous boosts, we conducted passive antibody transfer followed by challenge with SARS-CoV-2. We chose to carry out passive antibody transfer to fully assess the ability of the humoral response to protect against infection. We pooled sera from IFNAR^−/−^-CD46Ge mice boosted with wt or omicron-based MR-SARS-CoV-S6p312 and administered 150 µL of this serum into the peritoneum of K18-hACE mice, which express hACE2 under the epithelial cytokeratin promoter ^78^. As a mock-vaccination control, serum from IFNAR^−/−^-CD46Ge mice vaccinated twice with a MeV-MR empty vector was used. Animals were then challenged 2 h later by the intranasal route with 10^4^ pfu of 1) USA-WA1/2020 SARS-CoV-2 (Wuhan-like) or 2) omicron BA.1 virus. The mice were monitored for signs of clinical disease following infection, including daily weight changes. On day 5 post-infection, the mice were euthanized, and lung tissue nasal turbinates were collected to determine virus titers by plaque assay.

In mice challenged with USA-WA1/2020, those that were pretreated with vaccination serum showed no signs of weight loss. In contrast, weight loss was observed in the sham group mice starting at 4 days post-infection (dpi) (Figure 6D). Although substantial replication occurred in the lungs of passively immunized animals, both types of vaccination sera yielded similarly reduced lung viral titers (∼16-fold, Figure 6E). Only animals that received serum from wt MR-SARS-CoV-S6p312-vaccinated animals showed lower nasal viral titers (175-fold) than the empty MeV-MR control-vaccinated mice.

In mice challenged with BA.1 virus, we did not observe any body weight loss, and viral titers in the lung and nasal turbinates were ∼100-fold lower than those detected with USA-WA1/2020 (Figure 6E), as previously reported ^79^. Although SARS-CoV-2 was recovered in the lungs of all vaccinated mice after the challenge, no infectious virus was detected in the nasal turbinates. Notably, mice vaccinated with the omicron-based MR-SARS-CoV-2S6p312 vector showed ∼4-fold lower lung viral titers than wt construct- and mock-vaccinated animals. Thus, as suggested in a previous study ^24^, protection against BA.1 is modestly improved in animals treated with a BA1-based booster vaccine. We conclude from this experiment that in vitro antibody neutralization does not faithfully predict in vivo efficacy in a prophylactic setting.

### MR-SARS-CoV-S6p312 elicits SARS-CoV-2 spike-specific neutralizing antibody responses in the presence of pre-existing measles antibodies

A previously developed MeV/SARS-CoV-2 vaccine has failed to elicit nAb response in measles-immune individuals ^44, 45^. We postulated that our remodeled MeV based on the Moraten strain might be less vulnerable to this so-called “blunting effect’ if nAbs against the MeV coat are involved in dampening the heterologous humoral immune response to the transgene. To begin to address whether the previously observed pseudovirus-neutralizing responses could be hampered by measles immunity, we vaccinated IFNAR^−/−^-CD46Ge mice in the presence or absence of MeV-specific IgG. To this end, 400 mIU of MeV nAbs were administered three hours prior to vaccination with either the MR-CoV-S6p312 or the MeV Moraten vaccine, which was used as a control. Three weeks later, following another passive administration of MeV nAbs, the animals were administered a booster. We then assessed the nAb response against MeV (Moraten vaccine) or SARS-CoV-2 pseudoviruses, as well as T-cell immunity (Figure 7A). As we expected, MeV nAbs were not detected in mice vaccinated with MR-CoV-S6p312 due to the serologically distinct MeV coat ^46^. In contrast, naïve animals vaccinated with the homologous Moraten virus developed a mean MeV neutralization titers of 6,194 mIU/mL. However, when animals were passively immunized, the production of MeV nAbs was reduced at 136 mIU/mL (Figure 7B). These results indicate that we successfully mimicked the impact of pre-existing anti-MeV antibodies on the immunogenicity of the measles vaccine ^80, 81^.

**Figure 7.**
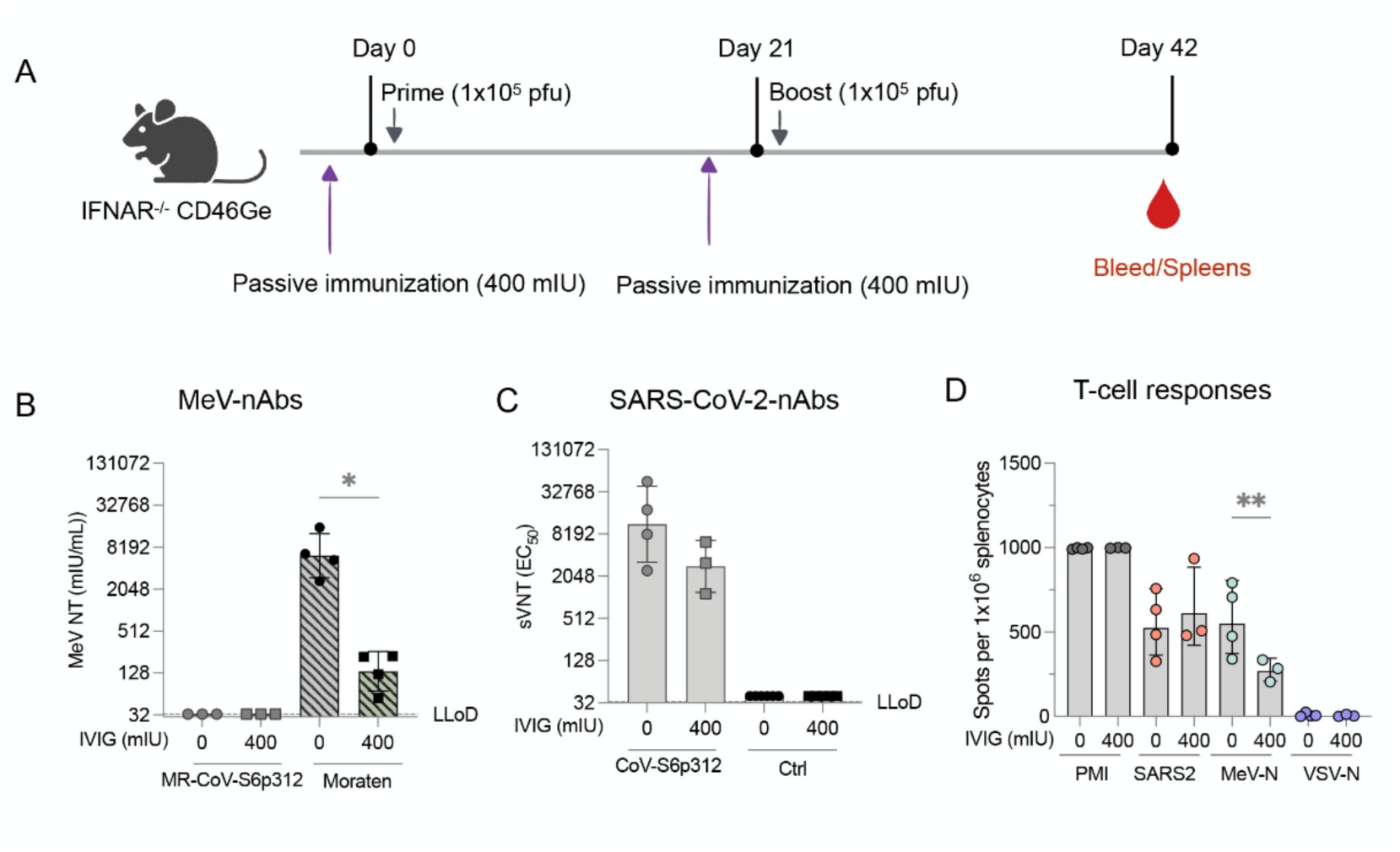
Pre-existing anti-measles virus antibodies do not blunt the anti-SARS-CoV-2 spike immune response elicited upon vaccination with MR-CoV-S6p312. **(A)** Study schematic of the experimental design. IFNAR^−/−^-CD46Ge mice were passively immunized on days 0 and 21 with 400 mIU of anti-measles virus neutralizing antibodies before each vaccination with a dose of 1x10^5^ pfu of MR-CoV-S6p312. MeV Moraten was used as a control for vaccination. Serum samples were collected three weeks after the second vaccination dose and tested for **(B)** MeV-neutralizing antibodies and **(C)** SARS-CoV-2 spike pseudovirus-neutralizing antibodies. **(D)** ELISPOT assays were performed on splenocytes collected three weeks after the second vaccination. Dots represent individual mice, and bars and error bars depict the geometric mean ± geometric standard deviation, respectively. Statistical analysis between groups was calculated by one-way ANOVA with Fisher’s LSD test (*, p<0.05; **, p<0.005).

Data obtained at the same time point were also collected to analyze the immune response against the SARS-CoV-2 spike protein generated in response to MR-CoV-S6p312. Similar levels of pseudovirus nAbs were present in naïve animals and animals with pre-existing anti-MeV antibodies. As we expected, pseudovirus nAbs were not detected after vaccination with the MeV Moraten vaccine. ELISPOT assays performed three weeks after the second dose revealed no significant differences (p>0.05) in the number of SARS-CoV-2 spike-specific IFNγ-producing cells in the animals vaccinated in the presence or absence of pre-existing anti-MeV antibodies. However, we did observe a significant decrease in the number of MeV-N-specific IFN-γ-producing cells. In conclusion, pre-existing MeV nAbs do not hamper the immunogenicity of our MeV-MR-vectored vaccine, since titers were comparable to those observed in naïve animals. Collectively, our results suggest that a MeV/SARS-CoV-2 vaccine candidate based on a remodeled MeV can be used as an effective strategy to elicit long-lasting nAb responses against SARS-CoV-2 virus in a measles-immune human population.

## Discussion

In this study, we sought to generate a remodeled live-attenuated measles vaccine capable of driving high nAb responses against the SARS-CoV-2 spike protein. The data presented here, show that antibodies alone resulting from a live-measles-vector COVID-19 vaccine candidate can protect against morbidity upon challenge with SARS-CoV-2. Our work provides direct and solid evidence that antigen optimization of the SARS-CoV-2 spike protein strongly enhances efficient induction of nAb production. These conclusions are based on the following evidence. First, we demonstrated that artificial trimerization of the SARS-CoV-2 spike protein is critical for the induction of an effective humoral (but not a cellular) immune response against the spike protein by measles-based COVID-19 vaccine candidates. Second, we showed that further multimerization of the SARS-CoV-2 spike by means of genetic fusion to H. pylori NAP further augments the magnitude of the nAb response. Third, we showed that boosting with an omicron-based COVID-19 vaccine candidate restores neutralizing activity against historical and contemporary SARS-CoV-2 variants. Finally, we provided strong evidence that pre-existing anti-MeV antibodies do not impact the immunogenicity of MeV-based COVID-19 vaccine candidates with epitope-modified H and F surface glycoproteins (MeV-MR). Overall, this experimental evidence strongly supports the further development of MeV-MR-based vaccine candidates expressing engineered SARS-CoV-2 spike protein to induce a protective immune response in measles-immune individuals.

The Comirnaty (BioNTech/Pfizer) and SpikeVax (Moderna) COVID-19 vaccines have saved millions of lives owing to their unprecedented speed of development and high degree of efficacy ^82^. During the first year of the pandemic, the two vaccines provided >95% efficacy against symptomatic infection, but since then, there has been substantial attrition in the ability of the current vaccines to reduce infection, likely due to the resulting selective pressure for immune-evasive variants. Although booster shots can “restore” nAb responses and protection against variants ^24, 83, 84^, the use of boosters will not end this pandemic, and there is an urgent need for next-generation vaccines. The live-attenuated MeV vaccine induces both humoral and cellular immune responses that can last the lifespan of an individual, possibly due to persistence in lymphoid tissues ^36, 85^. It is also one of the safest human vaccines ever developed, with outstanding safety records, especially in children <5 years old. These two features make the live-attenuated MeV vaccine an attractive platform as a vaccine vector against other pathogens. Although whether the longevity of the protection against measles is applicable to other diseases remains unknown, the high seroprevalence of MeV in the human population limits these studies, as exemplified recently by a phase I/II clinical trial of a measles-vectored SARS-CoV-2 vaccine candidate ^45^.

Here, we used our recently described MeV-MR to circumvent this blunting of vector immunogenicity ^46, 86^. We previously showed that, in contrast to the MeV vaccine strain, MeV-MR replicated in vivo in passively immunized animals ^46^. To investigate how to improve the immunogenicity of an MeV-MR-based COVID-19 vaccine candidate, we studied the antibody response elicited by various SARS-CoV-2 spike antigens in mice. In this approach, we focused on the full-length spike ectodomain. Even though the spike ectodomain (S1+S2) and the S2 subunit induced the production of similar levels of IgG antibodies that bound to the S1+S2 ectodomain and S2 subunit, we observed pseudovirus nAbs for only S1+S2-elicited antibodies. This observation confirms previous reports on the lack of immunogenicity of S2 in both mice and rabbits ^41, 87–89^. Although the lack of a quaternary structure for the purified spike ectodomain could have resulted in suboptimal responses, we also observed uniformly low nAb levels produced in response to the MeV-encoded full-length S construct, which contains the native transmembrane trimerization motif ^90^. Interestingly, two prior studies on MeV-based vaccine candidates expressing full-length spike achieved similarly low-to-absent nAb responses in cotton rats ^43^ and IFNAR^-/-^ KO CD46 mice ^42^, indicating the generalizability of our results across vectors and species. Although growing evidence indicates that multimeric antigens enhance immunogenicity more than soluble antigens ^63, 91^, it was previously unknown whether a trimerization motif could increase the immunogenicity of the full-length ectodomain of SARS-CoV-2 S protein ^43, 92, 93^. Building on work characterizing the prefusion conformation of the viral envelope and its antigenicity among different enveloped viruses ^12, 52, 94^, we designed a trimeric prefusion SARS-CoV-2 S protein by fusing the foldON trimerization motif in conjunction with the introduction of prefusion-stabilizing substitutions ^16^ (SARS-CoV-2S6p3). Our results showed that the protein was still not sufficiently immunogenic after a single injection of 5 µg of purified protein adjuvanted with alum. However, further multimerization by the addition of the NAP (S6p312) significantly enhanced the immunogenicity of the spike.

Since the immunogenicity of antigens could be adjuvant specific, we next studied the immunogenicity of different spike antigens in the context of MeV-vectored vaccine candidates. Although there was no major difference in antibody titers as measured by ELISA, only those constructs harboring an artificial trimerization domain and NAP induced the production of pseudovirus nAbs. Of note, an S6P construct induced the production of more uniformly high neutralizing titers than its S-2P counterpart when fused to NAP. The finding that a spike protein with six stabilizing proline substitutions resulted in higher neutralizing antibody titers than the version with two prolines is notable as revised structure-based designs of fusion glycoproteins do not always result in better immunogenicity despite improved protein expression and stability ^95^. Our observation on the superior immunogenicity of the HexaPro construct adds to a recent publication by Zhang et al. ^96^ during the preparation of this manuscript. On the contrary, a mRNA COVID-19 vaccine candidate based on the Hexapro variant and developed by Sanofi Pasteur failed to elicit nAbs in mice and non-human primates ^97^. Notably, the latter construct lacked a trimerization domain. Thus, a mRNA COVID-19 vaccine candidate based on a the HexaPro variant trimerized via an inserted peptide domain will likely be a promising vaccine candidate. Moreover, a clinical grade Newcastle disease virus/COVID-19 vaccine candidate has been generated based on the HexaPro variant in which the transmembrane domain and cytoplasmic tail of the spike has been replaced with those from the fusion protein of NDV (NDV-F) ^98^. Whether the NDV-F transmembrane region and cytoplasmic tail better trimerize the SARS-CoV-2 spike protein remains to be determined.

As observed with other COVID-19 vaccine candidates ^24, 83, 99, 100^, different SARS-CoV-2 variants partially or completely escaped the nAbs that were produced after a single vaccination with MeV/SARS-CoV-2. However, a second vaccination dose with /SARS-CoV-2 or MeV/SARS-CoV-2 BA.1 led to a substantial increase in Wuhan-specific nAb titers to peak values after the initial vaccination. Notably, BA.1-specific nAb titers were lower after a second dose of homologous MeV/SARS-CoV-2 than after a dose of the heterologous and matched MeV/SARS-CoV-2 BA.1, which was consistent with published data ^24, 101, 102^. When we compared protection during challenge studies in a small number of K18-hACE mice using establishment of passive immunity, there was no clear correlation between nAb titer and protection. Although BA.1 viral titers in the lower respiratory tract were reduced after boosting with MeV/CoV BA.1 but not MeV/CoV2, the difference did not reach statistical significance. This disparity might be explained by the Fc effector function of nonneutralizing antibodies ^103^, which were not quantified in this study. Alternatively, a larger animal experiment or pathological analysis of lung sections might produce further insights. Nonetheless, others have produced a modestly greater protective effect against BA.1 challenge with a BA.1-matched vaccine ^24, 83^, if any effect was observed.

Interestingly, there have been inconsistencies reported for several antigens depending on the use of different viral vector platforms. For instance, an adenovirus 26 (Ad26) or MVA virus expressing the unmodified full-length SARS-CoV-2 S induced strong neutralizing responses against SARS-CoV-2 ^104–106^, whereas we and others have observed a low-to-absent nAb response with MeVs expressing a similar construct ^42, 43^. This discrepancy might be explained by the fact that studies with MeV-based recombinant vaccine candidates were performed in type I IFN receptor knockout mice, which may have impaired induction of the humoral immune response ^42, 107, 108^. In agreement with this, we failed to elicit a robust nAb response in IFNAR-/-CD46Ge mice when we used a VSV(+G) SARS-CoV-2 vaccine candidate. Likewise, Mercado et al. ^109^ observed that an Ad26 expressing a secreted 2P-stabilized S antigen with deletion of the S1/S2 furin cleavage site and a foldON trimerization motif replacing its TM and CT domains (S.dTM.PP) elicited the production of nAb titers comparable to those produced using Ad26 expressing the unmodified full-length SARS-CoV-2 S (S and S.dCT). In contrast, a homologous construct in the context of MeV vaccination revealed that the trimeric and secreted S-2P form was clearly superior ^43^. We did not generate the MeV-based recombinant vaccine candidate that was reported by Lu et al. ^43^ (preS) because we observed poor immunogenicity of the protein even when we used a more stable HexaPro S variant; not all the animals exhibited seroconversion after a single dose. We noticed that, although Lu et al. vaccinated IFNAR-/-CD46Ge mice and cotton rats with this MeV-based COVID-19 vaccine candidate, they reported that binding antibodies were produced only in the former, whereas no binding or nAbs were observed in the latter. Furthermore, the binding antibody levels in mice were assessed only after a two-vaccination regimen with high doses of rMeV (ten times higher than what we used in this study). Since the authors performed a SARS-CoV-2 challenge experiment in actively vaccinated animals, the observed protective effect in that study could have been produced exclusively by cellular immune memory, as shown previously in the hamster model ^110^.

Another MeV-based COVID-19 vaccine candidate was recently reported by Frantz et al. ^41^. The authors used a prefusion-stabilized full-length S (S-2P) membrane-anchored antigen (SF-S2-dER) and showed that in the context of active measles vaccination, different rodent models were protected from SARS-CoV-2 challenge. Although no direct comparison for a MeV vector expressing SF-S2-dER or preS has been reported, these two constructs have been studied in the context of Ad26 infection ^109, 111^. Extrapolating similar immunogenicity across platforms, we expect SF-S2-dER to be 1.8-2.6-fold more potent than preS at inducing nAb responses. Therefore, even ignoring the fact that the HexaPro version induced the production of 39-fold higher nAb levels than S2-P, we expect the MeV-based COVID-19 candidate presented here to elicit the production of 4-6-fold more nAbs than the construct reported by Frantz et al. Nevertheless, one important advantage of using a soluble spike over a membrane-anchored spike is antigen camouflage/decoy to prevent interference by pre-existing antibodies. For instance, if a MeV/SARS-CoV-2 chimeric virus is used as the vaccine-vector, pre-existing antibodies against SARS-CoV-2 spike might interfere with vaccine efficacy because these antibodies can preclude vector infectivity, a prerequisite of any live-attenuated vaccine. This feature could be particularly important for the use of vectored vaccines as booster shots in previously SARS-CoV-2-infected or vaccinated individuals. Indeed, passive immunization of individuals with the monoclonal antibody bamlanivimab has been shown to lower the antibody titers elicited by either Comirnaty (BioNTech/Pfizer) or SpikeVax (Moderna) COVID-19 vaccines by up to twofold ^112^. Although modest, this twofold difference mimics those in the peak nAb titers produced by these two mRNA-based COVID-19 vaccines ^113^, with measurable clinical impact ^114^. Another important factor to consider is the potential tropism expansion of the new vector after the incorporation of viral glycoproteins. Adding to these concerns is the risk of vaccine-induced disease in certain populations or cross-species transmission as a result. These complicate the regulatory approval pathway since it has the potential to negate the already established safety profile of the vector, while also impacting existing manufacturing processes.

A number of vaccines have needed improvements to enhance the antigenicity and durability of the elicited immune response. Common bacterial vaccines, such as the Hib vaccine or the pneumococcal vaccine, use protein conjugates that extent the duration of the protection against disease exerted by the vaccines ^115, 116^. The use of self-assembling protein nanoparticles to multivalently display viral antigens has been another effective approach to enhancing the magnitude and breath of the nAb responses ^15^. Multivalent vaccines are more efficiently captured by antigen-presenting cells, which traffic and accumulate to lymph nodes to enhance immune processing ^117, 118^. However, class I viral fusion proteins have not always been displayed in the native trimeric form on the nanoparticle ^119–122^. Our results demonstrate that the addition of an exogenous trimerization motif to the SARS-CoV-2 spike protein is critical for the enhanced nAb responses when displayed on NAP. Powell et al., ^123^ recently reported the genetic fusion of ferritin to the Ebola glycoproteins (GP) had no effect on the elicited antibody responses. Perhaps, the lack of a trimeric Ebola GP precluded the benefit of GP multimerization, as we have described here for SARS-CoV-2. Hence, our present study can inform into the next-generation of vaccine candidates based on class-I viral antigens.

Unlike other nanoparticle vaccines for SARS-CoV-2 comprising computationally designed two-component nanoparticles, all the components of our nanoparticle vaccine are from nature and encoded in the measles vector; sidestepping in vitro assembly and purification of the nanoparticle complex. It is unlikely that NAP and the elicited antibodies are of obvious toxicity as recombinant NAP protein has been safely tested in phase I trials as a vaccine candidate for H. pylori ^124^. Moreover, a phase I clinical trial using an oncolytic measles virus expressing NAP to treat patients with metastatic breast cancer has been initiated. (ClinicalTrials.gov: NCT04521764) ^125^.

Our study has several limitations. (1) We did not formally examine surface display of the covalently linked NAP-tagged SARS-CoV-2 spike (SARs-CoV-2S6p312). Since unadjuvanted SARS-CoV-2S6p312 did not elicit the production of pseudovirus nAbs, the increased immunogenicity of the NAP-tagged spike cannot be attributed merely to a toll-like receptors agonistic effect ^59, 126^. Future studies using single-particle cryo-EM analysis are therefore needed to confirm the assumption that NAP spontaneously self-assembles and displays trimeric SARS-CoV-2 spikes. (2) Only a small cohort of K18-hACE 2 mice was used due to limited animal and antiserum availability from the MeV/CoV2 vaccinated IFNAR^−/−^-CD46Ge mice. Thus, follow-up experiments with larger cohorts are needed to expand upon and generalize our results. (3) We analyzed the immune response and protection against BA.1 omicron, but currently, BA.4/5 are now the dominant omicron-lineage viruses, and the Food and Drug Administration (FDA) recommends that newer vaccines contain these latest omicron variant sequences ^127^. We nonetheless do not anticipate our results to deviate significantly based on the premise that, in the context of breakthrough infection, prior BA.1 infection provides substantial protection against BA.5 ^128, 129^. Similarly, two bivalent mRNA vaccines including components against BA.1 or BA4/5 in addition to the parental mRNA-1273 showed in mice equivalent protective effect against BA.5 in the lungs ^130^. (4) Our analysis did not account for Fc-effector functions, which are important in controlling SARS-CoV-2 infection in the respiratory tract ^131–133^. Correlative studies on the therapeutic activity of purified IgG and their corresponding Fab fragment might provide some insights. (5) Mouse antisera were used for the passive immunization study due to the limited availability of human sera with sufficiently high nAb titers to significantly reduce the in vivo replication of the parental Moraten virus. Studies on nonhuman primates and ultimately humans will be required to test the translatability of our COVID-19 vaccine.

In summary, we have leveraged the robustness and versatility of the live-attenuated measles vaccine with the potential of nanoparticle platforms. Our results could lead to the next-generation human vaccines for coronaviruses and other important pathogens.

## Materials and Methods

### Cells and viruses

BHK cells (catalog number (Cat#) CCL-10, ATCC, Manassas, VA, USA) were maintained in Dulbecco’s modified Eagle’s medium (DMEM; Cat# SH30022.01, GE Healthcare Life, Pittsburgh, PA, USA) supplemented with 10% fetal bovine serum (FBS; Cat# 10437–028; Thermo Fisher Scientific, Waltham, MA, USA), 100 units/mL penicillin and 100 μg/mL streptomycin (Cat# 15140122, Thermo Fisher). Vero African green monkey kidney cells expressing a membrane-anchored single-chain variable fragment (scFv) specific for a hexahistidine peptide (6× HIS-tag) ^134^ were cultured in DMEM supplemented with 5% FBS. Cells were incubated at 37°C in 5% CO_2_ with saturating humidity. The Indiana strain-based VSV expressing SARS-CoV-2 spike in place of VSV-G and trans-complemented with VSV-G has been described elsewhere ^66^. The recombinant measles virus (rMeV) based on the Moraten vaccine strain expressing firefly luciferase has been described previously ^46^. SARS-CoV-2 virus stocks were grown in TMPRSS2-overexpressing Vero-E6 cells maintained in DMEM supplemented with 10% FBS, 100 unit/mL penicillin, 100 µg/mL streptomycin, 1% nonessential amino acids (NEAAs), 3 µg/mL puromycin and 100 µg/mL normocin. USA-WA1/2020 virus was obtained from BEI Resources (NR-52281), and omicron BA1 virus (hCoV-19/USA/NY-MSHSPSP-PV44476/2021, GISAID: EPI_ISL_7908052) was obtained from the Mount Sinai Pathogen Surveillance Program at the Icahn School of Medicine at Mount Sinai.

### Constructs and virus rescue

The codon-optimized gene encoding Wuhan-Hu-1 (GenBank MN908947.3) was used as the basis for all SARS-CoV-2 spike constructs. The beta variant of the SARS-CoV-2 spike protein (L18F, D80A, D215G, del242/243, R246I, K417N, E484K, N501Y, A701V) was synthesized in two fragments (GENEWIZ, South Plainfield, NJ, USA) and cloned into the pcDNA3.1+ expression vector (Cat# V79020, ThermoFisher Scientific, Waltham, MA, USA) using an InFusion HD Kit (Takara, Shinagawa, Tokyo, Japan). All the other variants were obtained from InvivoGene (Toulouse, France). Amino acid substitutions and deletions in the SARS-CoV-2 spike protein (shown in Figure 2) were introduced using standard molecular biology techniques^135^ and confirmed by Sanger sequencing (GENEWIZ). When indicated, a C-terminal thrombin cleavage site (LEVLFQGP), a “foldON” sequence (GYIPEAPRDGQAYVRKDGEWVLLSTFL) ^55^ and H. pylori NAP (GenBank accession no. WP_000846461) were also incorporated at the extreme C-terminus of the construct.

All SARS-CoV-2 spike constructs were inserted directly by InFusion cloning into the Mlu/AatII sites of the pSMART LC MeVvac2 (eGFP)P vector encoding MeV-HΔ8/CDV-F ^46^. The inserts were modified at the stop codon to ensure compliance with the paramyxovirus rule of six ^136^. Rescue of rMeV was carried out on cotransfected BHK cells as described previously ^137^.

### Viral infections and multistep growth curves

Viruses were propagated by infecting Vero cells at a multiplicity of infection (MOI) of 0.03 in viral vaccine production serum-free medium (VP-SFM, Cat#11681020, ThermoFisher Scientific) supplemented with 20 mM of L-glutamine (Cat# 25030081, Thermo Fisher). Virus titers were determined using Vero cells preseeded in a 96-well plate at 10,000 cells/well and infected with serial tenfold dilutions in Opti-MEM I reduced-serum medium (Cat# 31985070, Thermo Fisher). After a 90 min absorption period, the cells were replenished with viral growth medium (DMEM+5% FBS). The titer was visually determined 2-3 days post-infection using a microscope and calculated in terms of plaque-forming units. For virus growth analysis, Vero cells were preseeded in a 6-well plate at 400,000 cells/well and infected at an MOI of 0.03. After a 1.5h absorption-period, the inoculum was removed, the cells were washed three times with Dulbecco’s phosphate-buffered saline (DPBS; Cat# MT-21-031-CVRF, Mediatech, Inc., Manassas, VA, USA), and the medium was replaced with 1 mL of VP-SFM. At various time points after infection, the cell culture fluid and cell lysates were harvested, and the virus titers were determined as described above.

### Next-generation sequencing (NGS)

RNA from virus stocks was extracted with the QIAamp Viral RNA Mini Kit (Cat# 52904, QIAGEN, Hilden, Germany), and one-step cDNA synthesis was then performed with SuperScript IV RT Viral cDNA (Cat# 12594025, ThermoFisher Scientific, Waltham, MA, USA) using the following pair of primers: F1/R3583 (5’-ACC AAA CAA AGT TGG GTA AGG ATA G-3’/5’-CAT TCA TCC TTC CTG TCG CCT AG-3’), F3409/R5488 (5’-AGC AAA GTG ATT GCC TCC CAA G-3’/5’-ATA TGG CAG AGA CGT TCA CCT TG-3’), F5380/R9560 (5’-ACA CCC GAC GAC ACT CAA C-3’/5’-GAG TTC ACG GAT CTT CCT CGT TG-3’), and F9473/R15894 (5’-GGC CCA CTC TCA TAT TCC ATA TCC-3’/5’-ATA TGG CAG AGA CGT TCA CCT TG-3’). DNA fragments were gel-purified using a QIAquick gel extraction kit (Cat# 28704, QIAGEN), and amplicon sequencing was performed by the Center for Computational and Integrative Biology (CCIB) DNA Core Facility at Massachusetts General Hospital (Cambridge, MA). Illumina-compatible adapters with unique barcodes were ligated onto each sample during library construction. Libraries were pooled in equimolar concentrations for multiplexed sequencing on the Illumina MiSeq platform with 2x150 run parameters. Upon completion of the NGS run, the data were analyzed, demultiplexed, and subsequently entered into an automated de novo assembly pipeline, UltraCycler v1.0 (Brian Seed and Huajun Wang, unpublished).

### BN-PAGE

Purified proteins were mixed with NativePAGE sample buffer (ThermoFisher) and loaded into a NativePAGE 4-12% Bis-Tris gel (Thermo Fisher) according to the manufacturer’s instruction. The BN-PAGE gels were run for 2h at 150V and stained with Coomassie blue.

### Negative-stain TEM

The complex samplewas diluted and an aliquot (3 μL) was placed on a thin,carbon-coated 200-mesh copper grid that had been glow discharged. After 1±0.1 min, excess solution was blotted with filter paper. The grid was washed by briefly touching the surface of the grid with a drop (30 μL) of distilled water on parafilm and blotting dry with filter paper. This touching and blotting step was performed three times, each time with a clean drop of distilled water. Three drops of 0.7% (w/v) uranyl formate negative stain on parafilm were then applied successively, and excessive stain was removed by blotting in the same fashion. The grid was allowed to remain in contact with the last drop of stain with the sample side down for 1-3 min in dark before removal of excessive stain and air dried at22 ± 1.5 °C. reporImages were collected with Talos L120C with an electron dose of ∼40 e-/Å2 and a magnification of 57kx and 92kx that resulted in a pixel size of 0.246 nm and 0.152nm at the specimen plane, respectively. Images were collected with an 4k×4K Ceta CMOS.

### Western blot analysis

Cells grown on a 6-well plate were infected with various rMeVs at an MOI of 0.03. At 36–48 h post-infection, the supernatant was collected and filtered through a 0.45-μm-pore membrane. Additionally, the cells were lysed in mammalian protein extraction reagent (M-PER; Cat# 78503, Thermo Fisher Scientific) supplemented with Halt protease inhibitor cocktail (Cat# 877886, Thermo Fisher Scientific). The protein concentration was determined using a Pierce Coomassie Plus Assay Kit (Cat# 23236, Thermo Fisher), and 3 μg of cell lysate or ∼20 μL of supernatant was separated on a precast 12% or 4-12% Bis-Tris polyacrylamide gel before being transferred to a polyvinylidene fluoride (PVDF) membrane using an iBlot2 dry blotting system (Thermo Fisher Scientific). The blot was then probed with anti-SARS-CoV-2-spike RBD (GTX135385, GeneTex, Irvine, CA, USA), anti-SARS-CoV-2 spike (Cat# GTX632604, GeneTex), anti-MeV nucleocapsid (Cat# LS-C144599, LsBio, Seattle, WA, USA) and anti-high affinity (HA) peroxidase (Cat#12013819001, Millipore Sigma, St. Louis, MO, USA) and developed with a KwikQuant western blot detection kit using a KwikQuant Imager (Kindle Bioscience LLC, Greenwich CT, USA).

### Expression and purification of antibodies

Heavy and light chain amino acid sequences were downloaded from the CoV-AbDab database and synthesized as codon-optimized gBlock fragments (GENEWIZ). These antibodies were expressed using the Expi293 expression system kit (Cat# A14635, Thermo Fisher) with human IgG1 constant regions. The culture supernatant was collected and loaded at 4 mL/min on a 5 mL HiTrap Protein G column (Cytiva, Marlborough, MA, USA) equilibrated with 10 mM phosphate, pH 7, using a Bio-Rad NGC fast protein liquid chromatography (FPLC) system. The medium was tittered to pH 7 with 1 M monosodium phosphate before loading. The antibodies were eluted with 100 mM glycine, pH 2.7, and collected in tubes containing 1 M dibasic sodium phosphate to neutralize the pH. The eluate was concentrated to <4 mL with a 4 mL 50 kDa molecular weight cut-off (MWCO) Amicon ultracentrifuge filter, and the buffer was exchanged on a 10 mL Zeba desalting column (ThermoFisher) equilibrated in PBS. The final antibody concentration was determined using the protein extinction coefficient for IgG (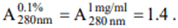).

### Generation of pseudovirus particles displaying SARS-CoV-2 spike and pseudovirus neutralization assay

Single-round pseudotyped lentivirus particles were produced by the cotransfection of HEK293T cells with the pHAGE-CMV-Luc2-IRES-ZsGreen-W (Cat# NR-52516, BEI), HDM-Hgpm2 (Cat# NR-52517, BEI), HDM-tat1b (Cat# NR-52518, BEI), pRC-CMV-Rev1b (Cat# NR-52519, BEI) and a SARS-CoV-2 spike plasmid as previously described ^48^. Virus-containing supernatants were harvested 72 h post-transfection, filtered using 0.45 μm syringe filters, aliquoted and stored at -80°C until further use. For neutralization assays, the virus was diluted to yield ∼ 50,000 relative light units (RLU)/well and incubated for 1 h at 37°C with 2-fold dilutions of heat-inactivated serum. Cells were then infected in quadruplicate and lysed 72 h later using the Bio-Glo luciferase assay system (Cat# GT7940, Promega, Madison, WI, USA) to measure luciferase activity. The percentage of neutralization was calculated based on the RLU measured using the virus-only control. The half-maximal effective concentration (EC_50_) titers were calculated using a log (agonist) versus normalized response (variable slope) nonlinear function in Prism 9 for macOS (GraphPad).

Alternatively, IMMUNO-CRON and IMUNO-COV v2.0 (Imanis Life Sciences) ^138^, which use a luciferase-encoding VSV displaying SARS-CoV-2 spike glycoproteins, were used to measure pseudovirus nAbs.

### Measles virus neutralization assay

A luciferase-based neutralization assay was used as previously reported ^46^. In brief, 2-fold serial solutions of serum samples were mixed with an equal volume of rMeV-Fluc and incubated for 1 h at 37°C. The virus-serum mix was subsequently added to Vero cells for 48 h before 50 nmoles of D-Luciferin (GoldBio, St Lous, MO, USA) was added to measure luminescence. The percentage of neutralization was calculated based on the RLU measured using the virus-only control and subsequently analyzed in Prism 9 to calculate the EC_50_ using a nonsigmoidal dose‒ response. EC_50_ values were converted into mIU/mL by using the third international standard for anti-measles serum (Cat# 97/648, National Institute for Biological Standards and Control).

### Mouse immunizations

All experimental procedures were carried out in accordance with US regulations and approved by the Mayo Clinic Institutional Animal Care and Biosafety Committee (IACUC). Male and female eight-to 19-week-old mice exhibiting deficient expression of type I interferon (IFN) receptor (IFNAR^-/-^) and transgenically expressing human CD46 (IFNAR^−/−^-CD46Ge)_ ^139^ were bred in-house under specific pathogen free conditions and regularly controlled by animal care takers and institutional veterinarians for general signs of well-being. Animals were maintained at a constant temperature of 22-25 °C, relative humidity of 40-70%, with a 12-h light/dark cycle and provided food and water ad libitum. For the experiments, animals were randomized for age-and sex-matched groups and no statistical consideration was taken. These animals were vaccinated intraperitoneally with 1x10^5^ plaque-forming units (pfu) of recombinant viruses or purified SARS-CoV-2 spike protein adjuvanted with aluminum hydroxide (alum, 2% alhydrogel adjuvant, Cat# vac-alu-250, InvivoGen, San Diego, CA, USA). A prime-boost vaccination regimen was used, and serum samples were collected before the vaccination booster was administered and at the end of the study. At this point, mice were euthanized, and splenocytes were harvested for study of the cellular immune responses. All serum samples were heat inactivated for 30 min at 56°C before humoral immune responses were assessed.

### Passive serum transfer

The transgenic K18-hACE2 mice (strain #:034860) were purchased from Jackson Laboratories and housed in a temperature-controlled vivarium with a 12 h day/night regime with water and food provided ad libitum. All experimental procedures were approved by the IACUC of the Icahn School of Medicine at Mount Sinai. For passive immunization, 150 µL of pooled serum was passively transferred by intraperitoneal injection 2 h before infection. The mice were infected intranasally with 10^4^ pf virus diluted in PBS, which was administered in 50 µL divided between both nostrils under mild ketamine/xylazine sedation (75 mg/kg ketamine; 7.5 mg/kg xylazine). Mice were monitored daily, and body weights were recorded. On day 3 or 5 post-infection, mice were euthanized via intraperitoneal injection of sodium pentobarbital (292.50 mg/kg). Lungs and nasal turbinates were isolated aseptically in 500 µL of PBS and homogenized for further use. Homogenates from lung and nasal turbinates were titrated to determine the virus load by plaque assay using TMPRSS2-expressing Vero cells as previously described ^140^.

### Recombinant SARS-CoV-2 antigens

Recombinant SARS-CoV-2 proteins produced in a baculovirus system were commercially obtained from Sino Biologicals, as follows: S1+S2 ectodomain, (Cat# 40589-V08B1) S1, (Cat# 40591-V08B1), RBD (Cat# 40592-V08B), S2, (Cat# 40590-V08B) and nucleocapsid (Cat# 40588-V08B). Trimeric SARS-CoV-2 spike and spike-H. pylori NAP proteins (SARS-CoV-2S6p3 and SARS-CoV-2S6p312, respectively) were produced upon transient expression in Expi293F cells (ThermoFisher). Clarified supernatants were purified by affinity chromatography using an anti-HA affinity matrix (Cat# 11 815 016 001, Millipore Sigma) pre-equilibrated with 20 mM Tris, 0.1 M NaCl, and 0.1 mM EDTA, pH 7.5 (equilibration buffer). The column was washed with equilibration buffer containing 0.05% Tween 20, and then elution was performed with 1 mg/mL HA synthetic peptide (Cat# 26184, Thermo Fisher) per the manufacturer’s instructions. Fractions containing the eluted proteins were combined, concentrated, and dialyzed against Dulbecco’s PBS (Cat# 25-508, Genesee Scientific) using a Pierce protein concentrator, 10 kDa molecular weight cutoff (MWCO, Cat# 88516, Thermo Fisher). The HA matrix was regenerated with 20 volumes of 0.1 M glycine, pH 2.0 (Cat# SC295018, Santa Cruz Biotechnology), and re-equilibrated before the next purification round. The protein concentration was determined using a Pierce 660 protein assay kit (Cat# 22662, Thermo Fisher). SARS-CoV-2S6p312 was purified after SARS-CoV-2S6p3 and stored at -80°C until use.

### Antigen binding enzyme-linked immunosorbent assay (ELISA)

IgG binding to SARS-CoV-2 or MeV antigens was measured by ELISA using clear flat-bottom immuno nonsterile 96-well plates (Cat# 442404, ThermoFisher Scientific) coated overnight at 4°C with 100 ng of recombinant SARS-CoV-2 proteins or 1 μg of MeV bulk antigen (Cat# BA102VS, Institut Virion\Serion GmbH, Würzburg, Germany) in 50 mM carbonate-bicarbonate buffer, pH 9.6. The plates were washed and blocked with 2% bovine serum albumin (BSA) in PBS for 2 h at room temperature (RT). The plates were washed again, incubated with serial dilutions of mouse serum and incubated for 1 h at 37°C. The plates were washed three times with PBS with 0.05% Tween 20 and then incubated for 1 h at RT with horseradish peroxidase (HRP)-conjugated anti-mouse IgG (1:5,000, Cat# 62-6520, ThermoFisher Scientific), IgG1 (1:5,000, Cat# 115-035-205, Jackson ImmunoResearch) or IgG2a (1:5,000, Cat# 115-035-206, Jackson ImmunoResearch) secondary antibodies. After the final wash, the plates were developed using 50 μL of 1-Step Ultra TMB (3,3’,5,5’-tetramethylbenzidine; Thermo Fisher Scientific), and the reaction was stopped with an equal volume of 2 M sulfuric acid before the optical density (OD) was read at 405 nm using an Infinite M200Pro microplate reader (Tecan). The endpoint titers of serum IgG responses were determined as the dilution in which the OD exceeding the average of the OD values plus three standard deviations of that of pooled negative serum samples was observed. Alternatively, anti-SARS-CoV-2 binding IgG levels were reported in units of μg/mL based on a standard curve that was generated using a SARS-CoV-2 spike nAb (Cat# 40595-MM57, Sino Biological).

### T-cell responses to viral antigens

IFN-γ enzyme-linked immunospot (ELISPOT) assays were carried out using mouse splenocytes to assess T-cell responses against MeV and SARS-CoV-2 peptides. Briefly, 5×10^5^ isolated splenocytes were cocultured with different stimuli in 200 μL of RPMI-10% FBS complete medium for 48 h on IFN-γ-coated plates (Cat# EL485, R&D systems, Minneapolis, USA). Fifteen-mer overlapping peptides from SARS-CoV-2 spike glycoprotein (Cat# PM-WCPV-S-1, JPT peptides, Berlin, Germany) and MeV-nucleoprotein (Genscript, NJ, USA) were used to stimulate splenocytes at 5 µg/mL. As a positive control, a phorbol myristate acetate (PMA)/ionomycin cell stimulation cocktail (Biolegend, San Diego, CA, USA) was used at 2.5 µL/mL, and as a negative control, splenocytes were stimulated with an equivalent DMSO concentration (0.8%). At 48 h post-incubation, the plates were developed in accordance with the manufacturer’s instructions. Developed IFN-γ spots were counted with an automated ELISPOT reader (CTL Analyzers LLC, USA). Each spot represented a single reactive IFN-γ–secreting T cell.

### Measurement of Th1/Th2 cytokines using ex vivo stimulation of splenocytes with antigen peptides

Frozen splenocytes were thawed and incubated with 50 µg/mL DNase1 (Cat# 10104159001, Roche) for 5 min at 37°C. The cells were then washed twice and resuspended in RPMI-1640 medium with 10% (vol./vol.) heat-inactivated FBS. Splenocytes (1×10^6^/well in 96-well plates) were stimulated for 24 h with 15-mer overlapping peptides from SARS-CoV-2 spike glycoprotein (Cat no# PM-WCPV-S-1, JPT Peptide Technologies GmbH) or VSV-N (Genscript) at a concentration of 2.5 µg/mL. Supernatants were collected, centrifuged at 1,800 RPM for 5 min and stored at -80 °C until analysis. Supernatants were then analyzed for the expression of IFN-γ, IL-6, IL-18, GM-CSF, IL-1β, IL12p70, IL-13, IL-2, IL-4, TNF-α and IL-5 cytokines using a mouse cytokine 11-plex antibody bead kit (Th1/Th2 Cytokine 11-Plex Mouse ProcartaPlex™ Panel, Cat No# EPX110-20820-901, Thermo Fisher). Preparation of samples, along with kit standards, detection antibody and streptavidin-phycoerythrin (PE), was carried out per the manufacturer’s instructions. Cytokine bead fluorescence intensity was measured using the Luminex 200 system (Luminex Corp., Austin, TX, USA), and data were quantitated with xPONENT® software.

### Statistical analysis

Statistical analyses were performed with GraphPad Prism version 9.1.0 for Mac OS 10.15.7. Significant differences among groups were determined as described in the figure legends.

## Acknowledgments

We thank the Mayo Clinic Proteomic Core and Dr. M. Cristine Charlesworth for antibody purification. The CCIB at Massachusetts General Hospital for the use of the CCIB DNA Core Facility (Cambridge, MA), which provides NGS services. We also thank Dr. Jesse Bloom for the 4^th^ generation lentivirus system and Dr. Vincent Munster for the SARS-CoV-2 spike expression plasmid used in the pseudovirus neutralization assays.

## Author contributions

Conceptualization: MÁM-A and SJR; Methodology: MÁM-A, RV, MS; Validation: MÁM-A, NP and RV; Formal analysis: MÁM-A, GS, RN and RV; Investigation; MÁM-A, RAN, BB, LZ, GS, PW, IM, RN; Resources: MÁM-A, RV, AG-S, MS, SJR; Writing-Original Draft; MÁM-A; Writing-Review and Editing: all authors; Visualization; MÁM-A; Supervision: MÁM-A, AG-S, MS, SJR; Project Administration: MÁM-A and MS; Funding acquisition: AG-S, MS, SJR.

## Data and materials availability

All data needed to evaluate the conclusions in the paper are present in the paper and/or the Supplementary Materials. Materials are available under a material transfer agreement.

