## Supplemental for "Surface-modified measles vaccines encoding oligomeric, fusion-stabilized SARS-CoV-2 spike glycoproteins bypass measles seropositivity, boosting neutralizing antibody responses to omicron and historical variants"

Miguel Á. Muñoz-Alía et al.

**This PDF file includes:**

Figs. S1 to S5

**Fig. S1.**

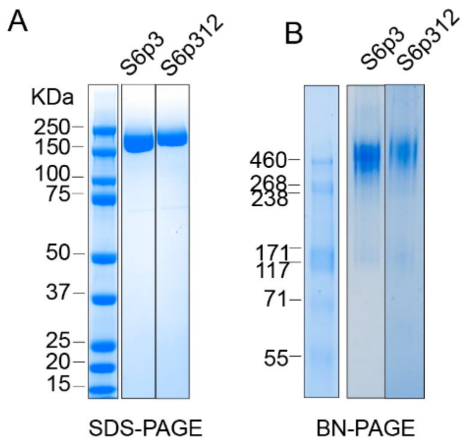


**Related to Figure 2.** (A) SDS–PAGE of the purified proteins. Protein (1 μg) was separated by SDS–PAGE (4-12% Bis-Tris gel) electrophoresis, followed by Coomassie staining. (B) BN-Native gel analysis of the purified proteins. Protein (1 μg) was separated by BN-Native electrophoresis (4-16% Bis-Tris), followed by Coomassie staining.

**Fig. S2.**

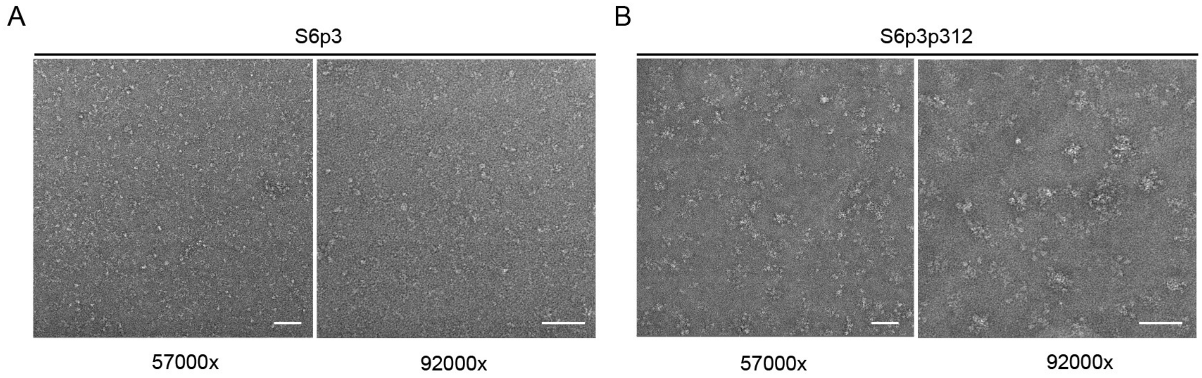


**Related to Figure 2.** Representative electron micrographs of negatively stained SARS-CoV-2S6p3 (A) and SARS-CoV-2S6p312 (B). Images were collected by Talos L120C 120 kV TEM with a magnification of 57kx and 92kx. Scales bars, 100nm.

**Fig. S3.**

| Virus | Gene | Nucleotide Mutation | Amino acid substitution |
|---|---|---|---|
| MR-SARS-CoV-S6pΔCT)P | ATU | t5565c | Y707H |
| | | t5579c | syn |
| | | t5626c | L727P |
| | | t5701c | L752P |
| | | t5712c | Y756H |
| | | t5721c/t5722c | F759P |
| | | t5738c | Syn |
| | | t5746c | L767P |
| | | t5811c | Y789H |
| | | t5825c | syn |
| | | t5827c | I794T |
| | | t5835c | F797L |
| | | t5844c | F800L |
| | | t5849c | syn |
| | | t5850c | F802L |
| | | t5860c | I805T |
| | | t5861c | syn |
| | | t5873c | syn |
| | | t5911c | L822P |
| | | t6108c | F888L |
| | | t6138c | F898L |
| | | del7140 | Frameshift |
| | H | c11335a | syn |
| MR-SARS-CoV- | ATU | c6175t | A890V |
| MR-SARS- | N | t398c | syn |
| | | t411c | syn |
| | | t467c | syn |

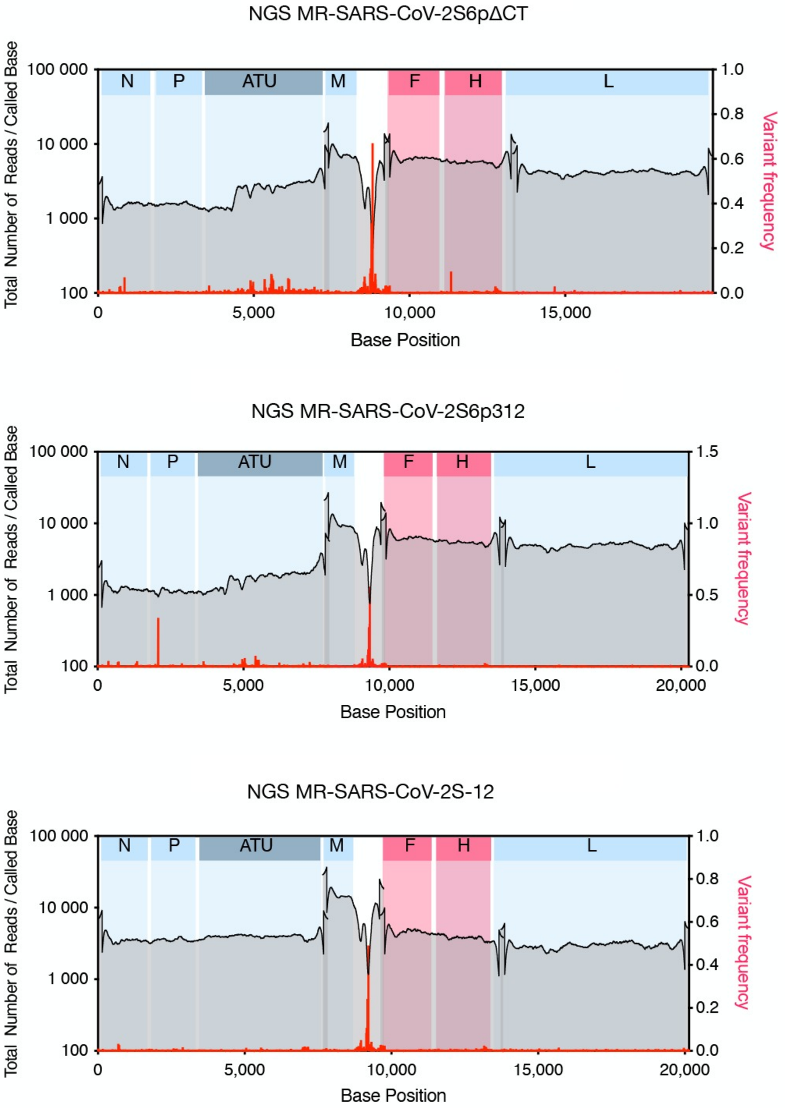


**Sequence divergence of rescued viruses from cloned cDNA.** Shown are rescued viruses in which nucleotide mutations were found. The genome coverage and allelic frequency of from Next-Generation Sequencing is also shown. Blue areas represent viral coding sequences, while white areas indicate intergenic regions and untranscribed terminal regions of the genome.

**FIG S4.**

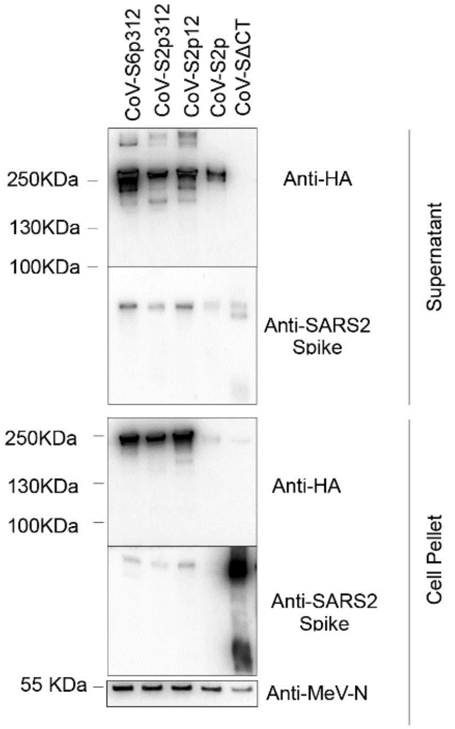


**Related to Figure 3**. Expression of SARS-CoV-2 spike-based constructs from the rMeV-MR vector. Vero cell lysates and supernatants were analyzed by western blotting two days after infection with the various spike-based measles vector constructs at an MOI of 0.03. The antibodies used for immunodetection as well as the molecular weight of a standard are indicated.

**FIG S5.**

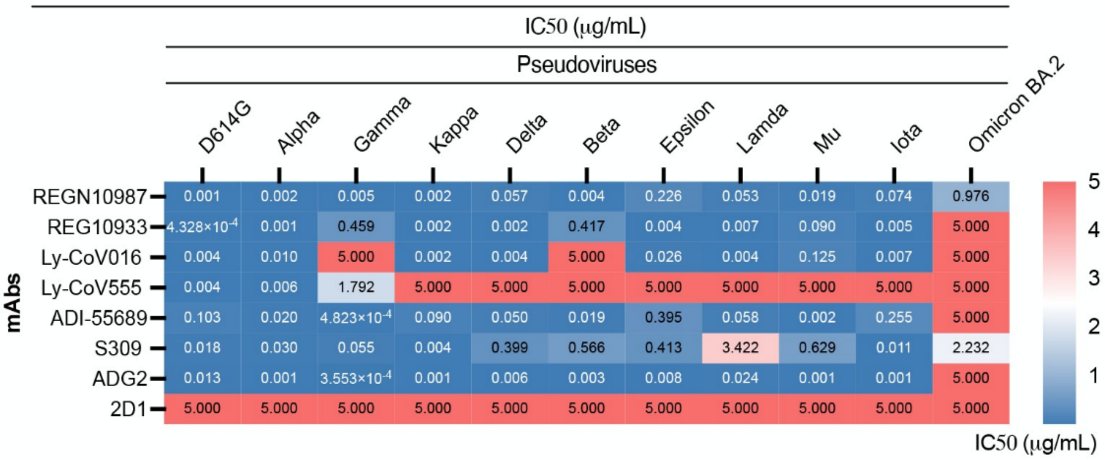


**Related to Figure 6. Pseudovirus neutralization assay of commercial mAbs against SARS-CoV-2 variants**. Half-maximal inhibitory concentration (IC50) values calculated from neutralization curves are plotted. 2D1 antibody was used as negative control.
